# KIF2C regulates synaptic plasticity and cognition by mediating dynamic microtubule invasion of dendritic spines

**DOI:** 10.1101/2021.08.05.455197

**Authors:** Rui Zheng, Yong-Lan Du, Xin-Tai Wang, Tai-Lin Liao, Zhe Zhang, Na Wang, Xiu-Mao Li, Ying Shen, Lei Shi, Jian-Hong Luo, Jun Xia, Ziyi Wang, Junyu Xu

**Author notes:** Corresponding author. Junyu Xu, Ziyi Wang, Jun Xia and Jian-Hong Luo. These authors contributed equally to this work (Rui Zheng, Yong-Lan Du).

## Abstract

Dynamic microtubules play a critical role in cell structure and function. In nervous system, microtubules specially extend into and out of synapses to regulate synaptic development and plasticity. However, the detailed polymerization especially the depolymerization mechanism that regulates dynamic microtubules in synapses is still unclear. In this study, we find that KIF2C, a dynamic microtubule depolymerization protein without known function in the nervous system, plays a vital role in the structural and functional plasticity of synapses and regulates cognitive function. Using RNAi knockdown and conditional knockout approaches, we showed that KIF2C regulates spine morphology and synaptic membrane expression of AMPA (α-amino-3-hydroxy-5-methyl-4-isoxazole-propionic acid) receptors. Moreover, KIF2C deficiency leads to impaired excitatory transmission, long-term potentiation, and altered cognitive behaviors in mice. Mechanistically, KIF2C regulates microtubule dynamics and microtubule invasion of spines in neurons by its microtubule depolymerization capability in a neuronal activity-dependent manner. This study explores a novel function of KIF2C in the nervous system and provides an important regulatory mechanism on how microtubule invasion of spines regulates synaptic plasticity and cognition behaviors.

## Introduction

Microtubules (MTs) are eukaryotic cytoskeletons that play an important role in cell function. In the central nervous system, MTs serve as a structural basis for trafficking material throughout the neuron and are the primary regulators of neuronal differentiation (*1*). The MTs dynamic instability, i.e. the growing or shrinking of MT plus end, has been detected in both shaft and dendritic protrusions (*2*) and is involved in spine formation and synaptic function (*3-6*). Inhibition of MT polymerization results in deficits in LTP and negatively affects memory formation or retention (*7, 8*). The abnormal MTs dynamics is associated with several neurodegenerative and psychiatric diseases, including Alzheimer’s disease, Parkinson’s disease, schizophrenia, and depression (*9*). During neuronal activities, microtubules can be regulated to extend from the dendritic shaft into spines and affect synaptic structure and function (*3-8*), which could be an important event for synaptic plasticity. However, regulatory mechanism underlying microtubule dynamics in or out from synapse upon neuronal activity is still unclear, nor its role in learning and memory.

The polymerization dynamics of MTs are essential for their biological functions and can be regulated by many MT-associated proteins (MAPs) (*10, 11*). The kinesin-13 family of kinesin superfamily proteins (KIFs) has a distinct function in depolymerizing MTs (*12*), and therefore is involved in many MT-dependent events (*13, 14*). KIF2C, also called mitotic centromere-associated kinesin (MCAK), is the best-characterized family member and is localized to the spindle poles, spindle midzone, and kinetochores in dividing cells. KIF2C utilizes MT depolymerase activity to regulate spindle formation and correct inappropriate MT attachments at kinetochores (*15-17*). KIF2C uses ATP hydrolysis for MT depolymerization from both ends and directly interacts with EBs to enhance its destabilizing ability (*18-22*). Particularly, KIF2C is associated with Alzheimer’s disease and suicidality psychiatric disorders. *In vitro* study showed that Oligomeric amyloid-β (Aβ) directly inhibits the MT-dependent ATPase activity of KIF2C and leads to abnormal mitotic spindles, which may cause chromosome mis-segregation and increased aneuploid neurons and thereby contributes to the development of Alzheimer’s disease (*23*). Clinical evidence shows that KIF2C is significantly decreased in suicidal ideation cohort and could be utilized as a reference for predicting suicidality (*24*). However, the *in vivo* physiological function of KIF2C in the central nervous system (CNS) remains to be elucidated.

In this study, we have found that KIF2C in the CNS is expressed in a brain-wide and synaptic localized pattern, which could be regulated by neuronal activity. Using a nervous system-specific KIF2C conditional knockout (cKO) mouse, we further confirmed its pivotal role in spine morphology, synaptic transmission, and synaptic plasticity. By overexpression of KIF2C wild-type (WT) and MT depolymerization-defective mutation, we have found that KIF2C regulates synaptic plasticity by mediating MT depolymerization and affecting MT synaptic invasion. Furthermore, KIF2C cKO mice exhibit deficiencies in multiple cognition-associated tasks, which could be rescued by overexpression of KIF2C WT but not the MT depolymerization-defective mutant. Overall, our study explores a novel function of KIF2C in neurons and synaptic plasticity, and reveals how MT dynamics in synapses upon neuronal activity contributes to cognition.

## Results

### KIF2C is highly expressed in the brain and at the synapse

Previous studies have showed that KIF2C plays an important role during cell division in mitotic cells (*25-27*); however, it is not clear whether KIF2C is also present in the nervous system. Western blot analyses showed that KIF2C was expressed in multiple regions of the brain (Figure 1A), and shows sustained high expression in the hippocampus throughout the postnatal stages (Figure 1B). To identify KIF2C abundance in neurons, western blot analyses were performed on whole-cell extracts of cultured hippocampal neurons (Figure 1C). The results showed that KIF2C was expressed in a development-related pattern in cultured neurons: detectable at day *in vitro* 3 (DIV 3) and significantly increased at DIV 10. Moreover, the subcellular fractionation analysis showed an enrichment of KIF2C in the postsynaptic density (PSD) fraction (Figure 1D). Immunofluorescent staining of cultured neurons revealed KIF2C expression as granular puncta of different sizes along dendrites overlapping with the postsynaptic marker PSD-95 and presynaptic protein Bassoon (Figure 1E). These results showed that KIF2C is abundantly expressed in CNS neurons and is enriched at postsynaptic density, suggesting a possible function in synapses.

**Figure 1.**
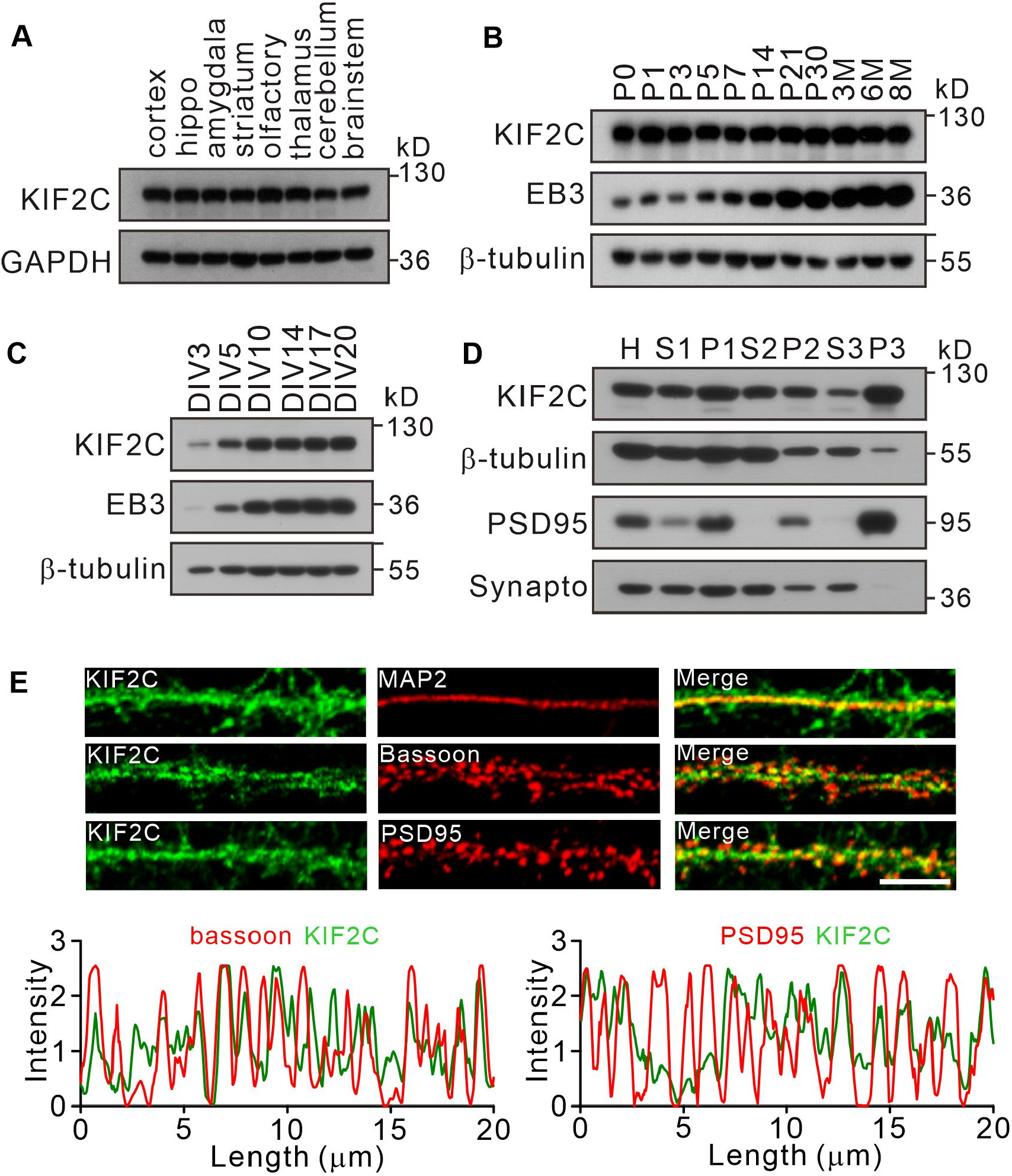
Expression of KIF2C in the central nervous system. (**A**) Expression of KIF2C in different mouse brain regions by immunoblotting. (**B**) Expression of KIF2C and EB3 in mouse hippocampus at different day *in vitro*. (**C**) Developmental expression patterns of KIF2C and EB3 in cultured hippocampal neurons. (**D**) Subcellular distribution of KIF2C in mouse hippocampus. H, homogenate; S1, low-speed supernatant; P1, nuclei; S2, microsomal fraction; P2, synaptosomal fraction; S3, presynaptosomal fraction; P3, postsynaptic density and synapto, synaptophysin. (**E**) Representative images of hippocampal neurons double-labeled with antibodies against KIF2C, PSD95 (postsynaptic marker) and Bassoon (presynaptic marker). Scale bar, 5 μm.

### KIF2C knockdown affects spine morphology, synaptic transmission and long-term potentiation *in vitro* and *in vivo*

To explore the function of KIF2C in synapses, KIF2C was knocked down in cultured hippocampal neurons using viral-mediated KIF2C shRNA (shKIF2C) (Figure 2A) at DIV 7–10 and the spine morphology was examined at DIV 16–18. The results showed that KIF2C knockdown did not affect spine density (Figure 2B). Electrophysiological recordings also showed similar frequency and amplitude of miniature excitatory synaptic currents (mEPSCs) in control and knockdown neurons (Figure S1A), suggesting that acute knockdown of KIF2C in cultured neurons does not affect basal synaptic transmission. Thereafter, we investigated whether KIF2C regulates spine morphology in an activity-dependent manner. The induction of long-term potentiation (LTP) results in changes in spine structure(*28-31*). To test whether KIF2C is involved in spine structural plasticity, we further conducted a glycine-induced chemical LTP (cLTP) assay to produce synaptic enhancements that mimic LTP(*32*) in KIF2C knockdown hippocampal neurons. Spine density did not change after cLTP stimulation (Figure 2C, D). However, the spine head width significantly increased in control hippocampal neurons, whereas it was not affected in the KIF2C knockdown neurons after cLTP stimulation (Figure 2E), indicating that KIF2C is essential for spine structural plasticity. In addition, both subcellular fractionation and immunocytochemical results showed that cLTP triggered the translocation of KIF2C from synaptic protrusions to the dendritic shaft (Figure 2F–I). Together, these data indicate that KIF2C might be removed from postsynaptic sites by neuronal activity, which may contribute to synaptic plasticity.

**Figure 2.**
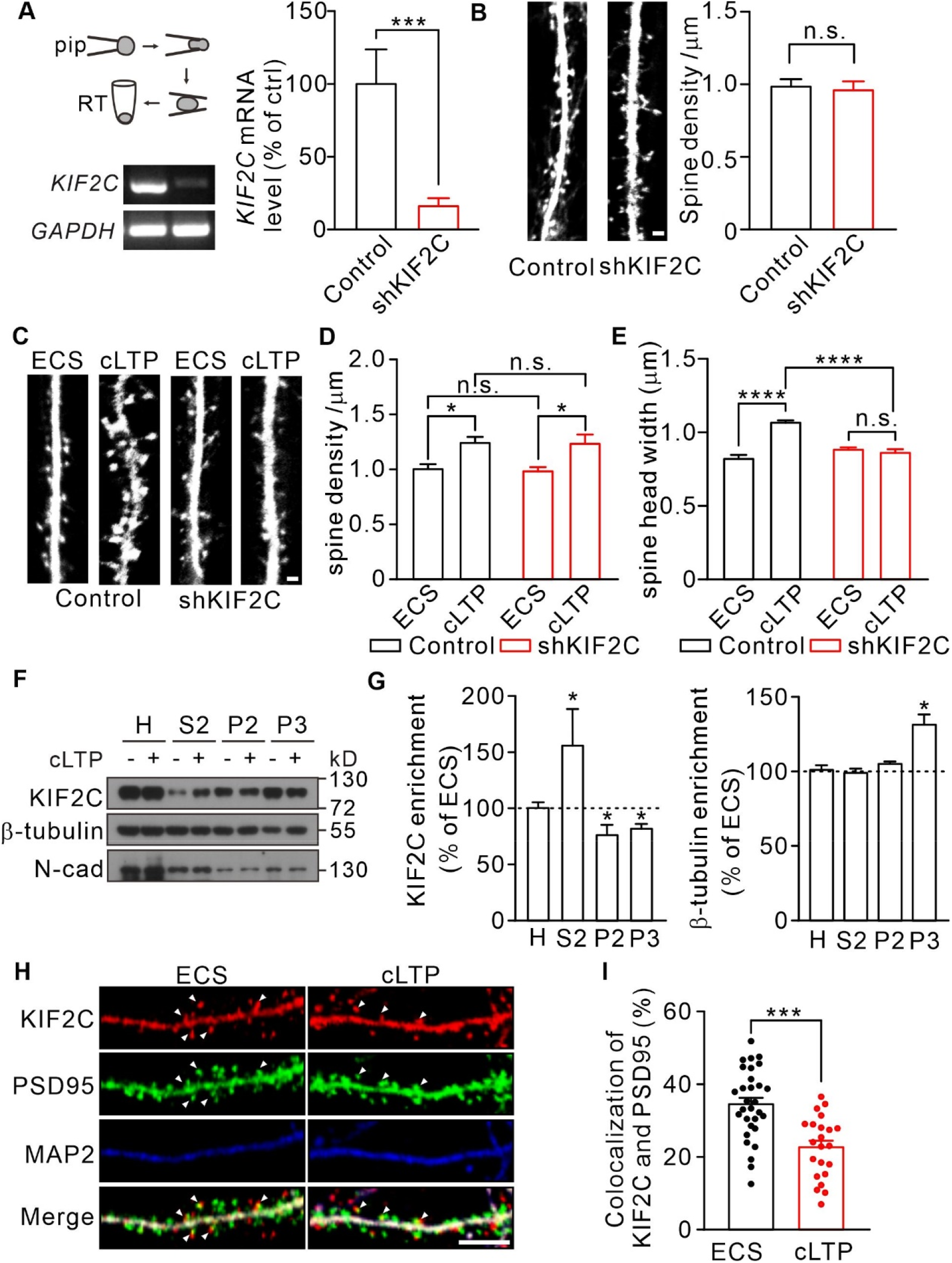
KIF2C displays translocation after cLTP and is required for synaptic structure plasticity. (**A**) Electrophoresis of *KIF2C* and *GAPDH* amplicons from individual control and shKIF2C hippocampal neurons. Histograms show percentage changes of *KIF2C* mRNA levels (% of control, \*\*\**p* < 0.001, *n* = 3 experimental repeats). (**B**) Cultured hippocampal neurons treated with control shRNA and shKIF2C at DIV 7-10, and captured at DIV 18-20. Scale bar, 1 μm. Right, Spine number per 1 μm, *n* = 20 neurons from 3 independent culture. (**C-E**) Cultured hippocampal neurons treated with control shRNA and shKIF2C at DIV10, and incubated with ECS or glycine (200 μM) for 10 min at DIV16-18. Scale bar, 1 μm. Spine density (**D**) and spine head width (**E**) were calculated. Data are mean ± SEM. Spine density, *n* = 17 neurons from 3 independent culture. Spine head width, *n* = 24 neurons from 3 independent culture. n.s., *p* >0.05, \**p* < 0.05, \*\*\*\**p* < 0.0001; Student’s *t*-test for spine density. One way-ANOVA for spine head width. (**F**) Subcellular fraction of KIF2C in cultured hippocampus neurons after 10 min of glycine treatment. N-cadherin (N-cad) was served as negative control. H, Homogenate; S2, microsomal fraction; P2, synaptosomal fraction and P3, postsynaptic density from ECS and cLTP treated neurons. (**G**) Histograms show percentage changes of KIF2C and β-tubulin in neurons with or without cLTP treatment. \**p* <0.05; Student’s *t*-test, *n* = 3 experimental repeats. (**H**) Representative images of co-localization of KIF2C (red), PSD95 (green) and MAP2 (blue). Scale bar, 5 μm. (**I)** Histograms show percentage changes of co-localization of KIF2C and PSD95. n =20 neurons from 3 cultures. \*\*\**p* < 0.001; Student’s *t*-test.

To further determine the effect of KIF2C on synaptic function, KIF2C was knocked down in hippocampal CA1 neurons *in vivo* and the electrophysiological properties in acute slices were measured. Mice were injected with either AAV-scramble shRNA (control) or AAV-KIF2C shRNA (shKIF2C) in the CA1 region (Figure 3A), and mEPSCs were recorded. Compared to the control, KIF2C knockdown neurons had enhanced amplitude but similar frequency of mEPSCs (Figure 3B–D). This discrepancy between cultured neurons and brain slices may result from activity-dependent synaptic scaling in free-behaving mice after viral injection. In addition, the excitatory postsynaptic current ratio of N-methyl-D-aspartic acid receptor (NMDAR)/α-amino-3-hydroxy-5-methyl-4-isoxazolepropionic acid receptors (AMPARs) were decreased in knockdown hippocampal slices (Figure 3E). Furthermore, we investigated whether KIF2C regulates presynaptic function retrogradely by comparing paired-pulse ratios (PPRs). KIF2C knockdown neurons exhibited comparable PPRs to the control, suggesting that KIF2C had a minimal effect on contacting presynaptic function (Figure 3F). To test whether LTP is affected in KIF2C knockdown neurons, we measured whole-cell LTP in CA1 pyramidal neurons by stimulating Schaffer collaterals. The magnitude of LTP was significantly attenuated within 5 min in knockdown hippocampal slices (Figure 3G, H). These results suggest that KIF2C is necessary for synaptic plasticity in hippocampal neurons, and its insufficiency may affect the basal function of NMDARs or AMPARs in the hippocampus.

**Figure 3.**
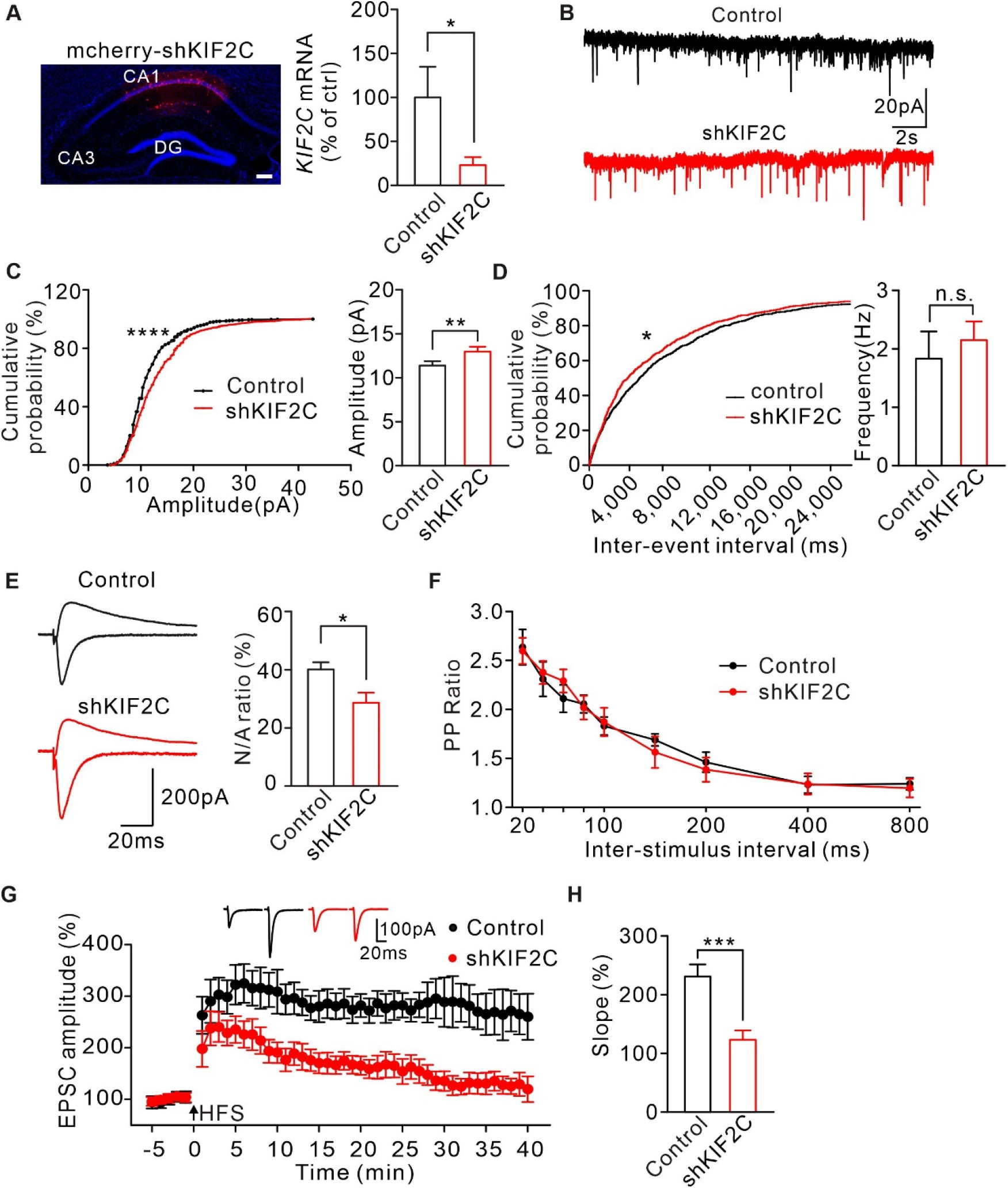
Knockdown KIF2C impairs synaptic transmission and plasticity. (**A**) Diagram of mcherry-shKIF2C injection, and coronal section image of the CA1 region. Scale bar, 100 μm. Histograms show percentage changes of KIF2C mRNA levels (% of control, \**p* < 0.05, n = 10 cells). (**B-D**) mEPSCs recorded from control (*n* = 10) and shKIF2C (*n* = 10) CA1 pyramidal neurons. n.s., *p* > 0.05, \*\**p* < 0.01, Student’s t-test; For the cumulative probability curves, \**p* < 0.05, \*\*\**\*p* < 0.0001, Kolmogorov-Smirnov test. (**E**) AMPAR and NMDAR EPSCs in CA1 pyramidal neurons. Left, sample sweeps illustrating NMDAR-EPSCs recorded at +40 mV (top trace) and AMPAR-EPSCs recorded at −70 mV (bottom trace). Right, histograms show ratio of NMDAR-EPSCs amplitude / AMPAR-EPSCs amplitude. \**p* = 0.0276. (**F**) Paired-pulse stimuli evoked EPSPs with different interval. (**G**)Time course of percentage changes of EPSCs amplitudes in control and shKIF2C CA1 pyramidal neurons. Each data point represents the average of 16 successive EPSCs evoked. (**H**) Histogram shows fEPSC slope of control and shKIF2C neurons (35-40 min). \*\*\**p* < 0.001; Student’s *t*-test.

### KIF2C knockout affects spine morphology, synaptic transmission, and long-term potentiation *in vivo*

To further investigate the functions of KIF2C *in vivo*, we generated cKO mice using the CRISPR/Cas9 system (Figure 4A). To obtain cKO mice, KIF2C^flox/flox^ mice were mated with Nestin-Cre transgenic mice, which resulted in KIF2C deletion only in the nestin cell lineage (Figure S2A). The knockout efficiency was confirmed using a hippocampal tissue RT-PCR assay (Figure S2B). There were no significant differences in body weight or brain size between Nestin-Cre;KIF2C^F/F^ (cKO) and KIF2C^F/F^ (WT) mice (Figure S2C). In addition, using Nissl staining, we found that the thickness of hippocampal CA1 and CA3 regions did not change significantly in cKO mice (Figure 4B). Further examination of neuronal morphology of CA1 pyramidal cells by Golgi staining and Sholl analysis showed no significant differences in dendritic complexity or spine density between KIF2C cKO mice and WT mice (Figure 4C–E). However, the density of mushroom-type spines decreased in basal and apical dendrites of hippocampal neurons in cKO mice, whereas the density of filopodia-type spines increased (Figure 4F). Furthermore, electron microscopy analysis revealed that the length and thickness of the PSD fraction were remarkably reduced in the hippocampus of cKO mice (Figure 4G, H). Mushroom spines are commonly considered mature spines(*33, 34*); therefore, these results indicate impaired spine maturation in the hippocampal CA1 region of cKO mice.

**Figure 4.**
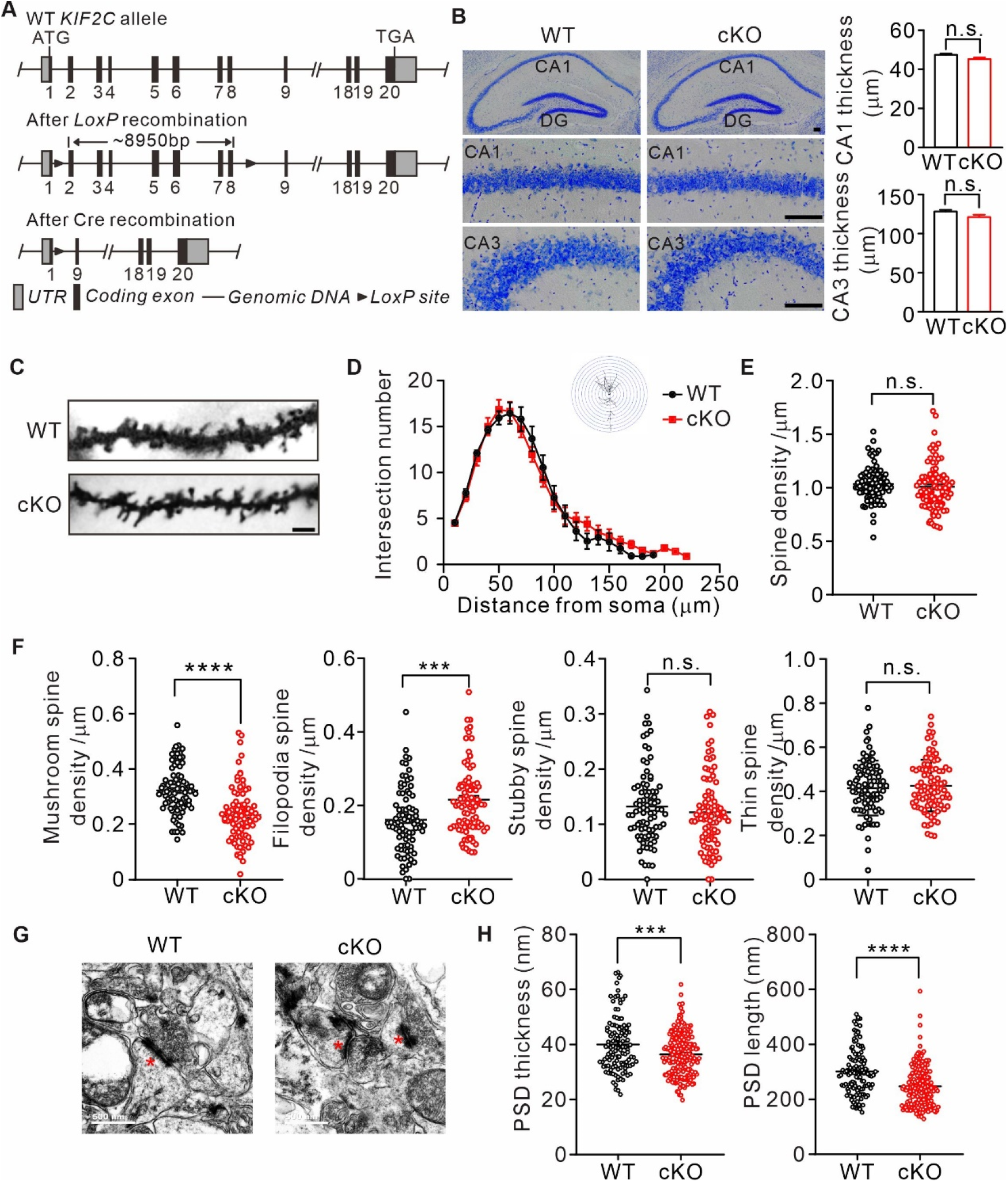
Abnormal spine formation in KIF2C conditional knockout mice. (**A**) Targeting strategy to generate KIF2C^flox/flox^ mice. sgRNA direct Cas9 endonuclease cleavage in intron 1-2 and 8-9 and create a double -strand break. (**B**) Nissl staining of adult mice brain coronal sections and magnified images of hippocampal CA1 and CA3 regions. Scale bar, 50 μm. (**C-F**) Golgi staining of WT and cKO hippocampus CA1 pyramidal neuron. *n* = 4 mice per genotypes; Scale bar, 2 μm. (**D**) Sholl analysis of dendritic arborization. The number of intersections related to distance from soma was quantified. *p* = 0.2824, two-way ANOVA RM and *post hoc* comparisons. (**E**) Histograms show spine number per 1 μm of WT and cKO CA1 pyramidal neuron; *p* = 0.3809. (**F**) The density of four types of spines in WT and cKO CA1 pyramidal neuron. Mushroom spine density was 0.32 ± 0.009 μm^-1^ (WT) and 0.23 ± 0.01 μm^-1^ (cKO) (\*\*\*\**p* < 0.0001). Filopodia spine density was 0.16 ± 0.009 μm^-1^ (WT) and 0.22 ± 0.01 μm^-1^ (cKO) (\*\*\**p* < 0.001). Stubby spine density was 0.12 ± 0.009 μm^-1^ (WT) and 0.13 ± 0.01 μm^-1^ (cKO) (*p* = 0.3104). Thin spine density was 0.41 ± 0.009 μm^-1^ (WT) and 0.43 ± 0.01 μm^-1^ (cKO) (*p* = 0.5511). (**G**) Transmission EM of synapse in hippocampal CA1 region from WT and KIF2C cKO mice (*n* = 3 mice per genotype) at 6-8 weeks. Scale bar, 500 nm. (**H**) Quantification of PSD thickness and length. PSD thickness: 40.02 ± 0.93 nm (WT) and 36.37 ± 0.58 nm (cKO). PSD length: 301.4 ± 6.8 nm (WT) and 247.7 ± 5.4 nm (cKO). *n* = 3 mice per genotype. \*\*\**p* < 0.001; \*\*\*\**p* < 0.0001; Student’s *t*-test.

To determine whether such impaired spine maturation may affect synaptic transmission and synaptic plasticity in cKO mice, mEPSCs were first examined in CA1 hippocampal pyramidal neurons. Consistent with the shRNA knockdown experiments in CA1, the amplitude of mEPSCs increased in cKO mice compared to that in WT controls (Figure 5A, B); whereas the frequency of mEPSCs was similar between the two groups (Figure 5C). The increased mEPSC amplitude was not due to altered presynaptic activity because PPRs were not affected in cKO neurons (Figure 5D), indicating a postsynaptic influence in mEPSC property changes. Therefore, we examined the NMDA/AMPA ratio of the CA1 neurons and found a decreased NMDA/AMPA ratio in cKO mice (Figure 5E). In addition, we performed cell-surface biotinylation experiments to assess possible changes in the surface expression of these two receptors. The levels of surface AMPA receptors significantly increased in the hippocampus of cKO mice, with NMDA receptors showing an increasing trend (Figure 5F), which may explain the increased mEPSC amplitude. We also compared long-term plasticity in the hippocampal CA1 region between WT and cKO mice and found that the LTP magnitude was significantly attenuated within 1 h in cKO hippocampal slices (Figure 5G, H), while long-term depression was not altered (Figure 5I, J), suggesting that KIF2C plays a critical role in LTP induction except for LTD.

**Figure 5.**
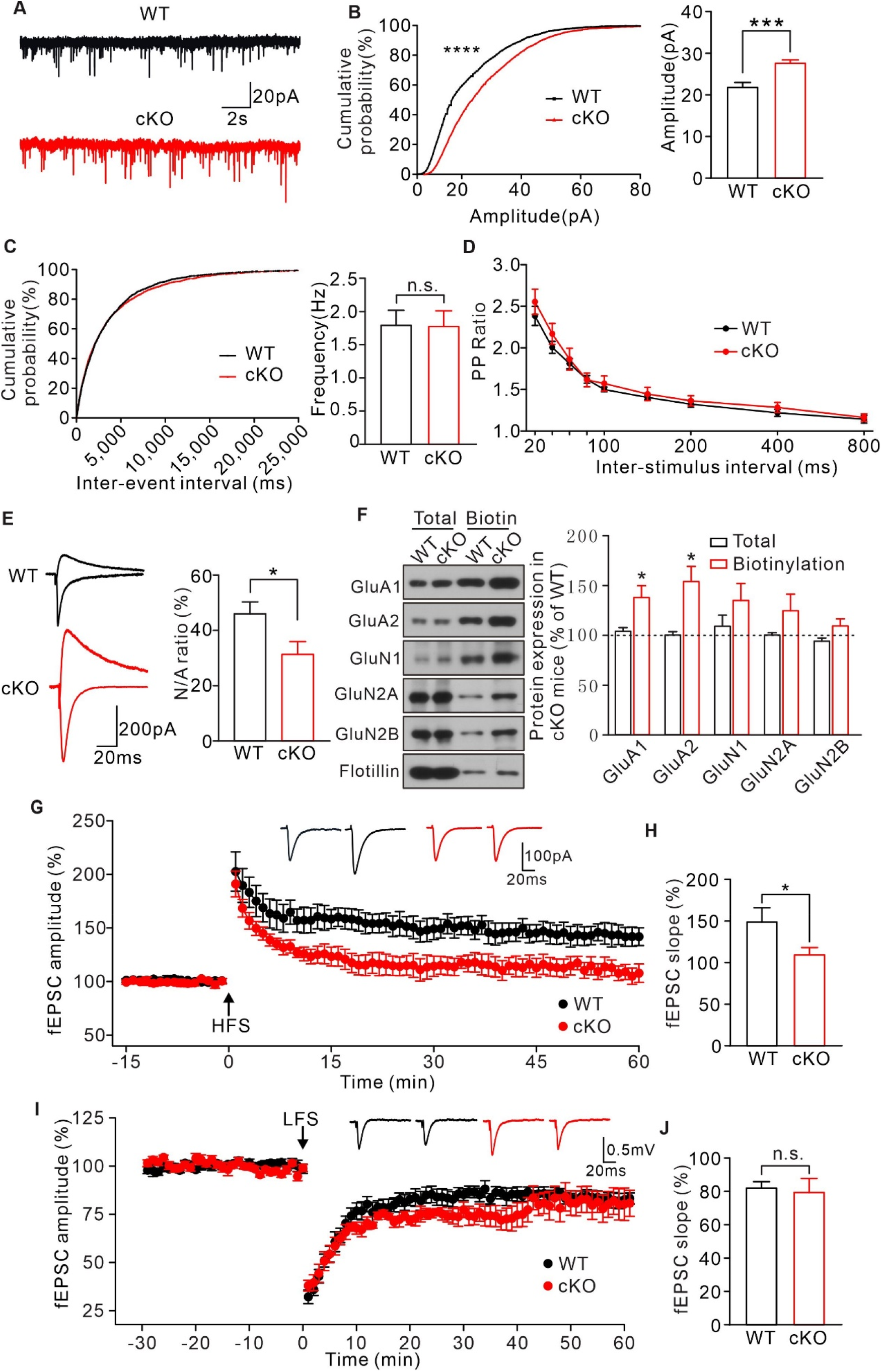
KIF2C deficiency impairs synaptic transmission and plasticity. (**A-C**) mEPSCs recorded from WT (*n* = 21) and cKO (*n* = 21) CA1 pyramidal neurons. \*\*\**p* < 0.001, Student’s *t*-test; For the cumulative probability curves, n.s. *p* > 0.05, \*\*\*\**p* < 0.0001, Kolmogorov-Smirnov test. (**D**) Paired-pulse stimuli evoked EPSPs with different interval. (**E**) AMPAR and NMDAR EPSCs in CA1 pyramidal neurons. Left, sample sweeps illustrating NMDAR-EPSCs recorded at +40 mV (top trace) and AMPAR-EPSCs recorded at −70 mV (bottom trace). Right, Histograms show ratio of NMDAR-EPSCs amplitude / AMPAR-EPSCs amplitude. *p* = 0.0322. (**F**) Cell-surface biotinylation shows a significant increase in GluA1 and GluA2 surface expression in cKO hippocampus compared with WT. Histograms show percent change of GluA1, GluA2, GluN1, GluN2A, GluN2B in total and biotinlation fraction. Flotillin was the internal control. \**p* < 0.05. (**G**) Time course of percentage changes of fEPSCs amplitudes in control and cKO CA1 slices. WT, *n* = 9 slices from 7 mice; cKO, *n* = 7 slices from 6 mice. (**H**) Histogram shows fEPSC slope of WT and cKO neurons (55-60 min). \**p* < 0.05, Student’s *t*-test. (**I**) Time course of percentage changes of fEPSCs amplitudes before (baseline) and after LTD stimulation in control and cKO CA1 slices. WT, *n* = 7 slices from 5 mice; cKO, *n* = 8 slices from 6 mice. (**J**) Histogram shows fEPSC slope of WT and cKO neurons (55-60 min). *p* = 0.7643, Student’s *t*-test.

### KIF2C regulates synaptic transmission and plasticity by mediating dynamic microtubule invasion of dendritic spines

Extensive studies on KIF2C in dividing cells have shown that KIF2C regulates microtubule dynamics by mediating microtubule depolymerization (*15-17*). However, dynamic microtubules are reported to be involved in regulating dendritic spine morphology and synaptic plasticity (*35*). Therefore, KIF2C may regulate synaptic plasticity through dynamic microtubules. To further explore this, we transfected EB3-tdtomato in DIV 7–10 WT or cKO hippocampal neurons to mark the MT plus end and observed the MT dynamics in mature neurons at DIV 18–20 using live-cell imaging. The results showed that the speed of EB3 was faster in cKO dendrites (WT: 8.9 ± 0.8 μm / min, cKO: 12.4 ± 0.5 μm / min) (Figure 6A, B), indicating an increased rate of MT polymerization in cKO mice. Using the plusTipTracker software, which is designed to track MT growth parameters(*36*), we analyzed the polymerization properties of the EB3-labeled MT plus ends in dendrites. Depletion of KIF2C significantly increased MT plus-end growth speed (WT: 18.1 ± 0.7 μm / min, cKO: 32.3 ± 1.4 μm / min); however, it decreased the growth lifetime (WT: 10.2 ± 0.4s, cKO: 6.4 ± 0.1s) (Figure 6B). MT plus end growth length was not significantly altered (WT: 2.7 ± 0.1 μm, cKO: 2.9 ± 0.1 μm) (Figure 6B). These results suggest that KIF2C regulates MT depolymerization in hippocampal neurons and that MT dynamics become disordered with KIF2C deficiency.

**Figure 6.**
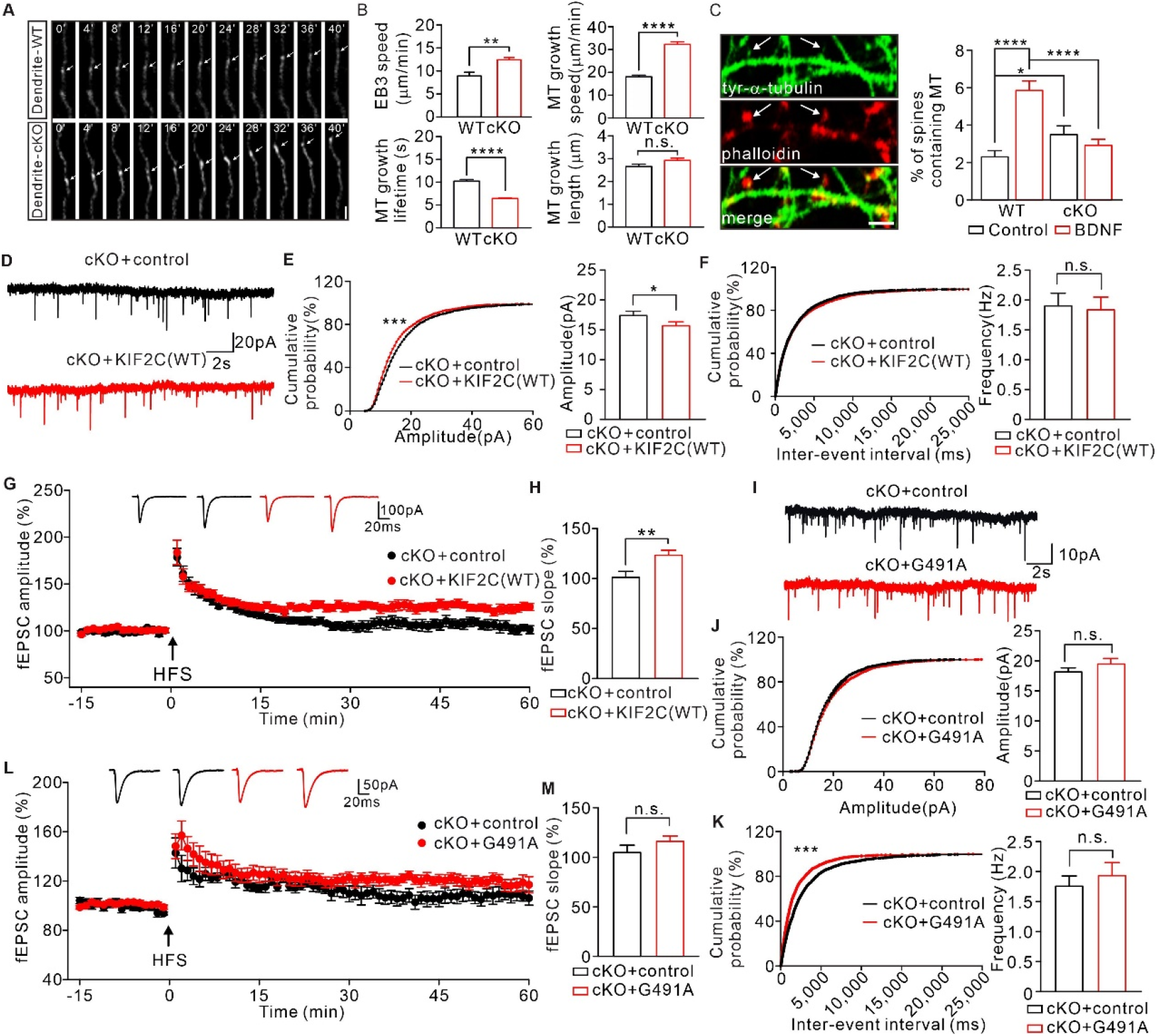
Abnormal MT dynamics cause deficit in synaptic transmission and plasticity. (**A**) Time-lapse recordings of EB3-tdtomato-infected neurons showing EB3-tdtomato comets in dendrite of WT and cKO hippocampus neuron. Subsequent images show low-pass-filtered time series in row. Scale bar is 1 μm, time in seconds. (**B**) Analysis of the EB3 comet dynamics in dendrites of WT and cKO hippocampus neuron. *n* = 108 from 7 independent experiments. Histograms show EB3 speed velocity (\*\**p* < 0.01), MT growth speed (\*\*\*\**p* < 0.0001), MT growth lifetime (\*\*\*\**p* < 0.0001) and MT growth length (*p* = 0.094). (**C**) Tyrosinated α-tubulin antibody-labeled MTs (white arrows) are detected in a small percentage of phalloidin (F-actin) -labeled spines under basal conditions (DIV 20). Quantification of the number of spines containing MTs before and after BDNF treatment. Scale bar, 5 μm. 2.4 ± 0.32% (WT), 5.9 ± 0.5% (WT + BDNF), 3.6 ± 0.44% (cKO), 2.9 ± 0.29% (cKO + BDNF). \**p* < 0.05, \*\*\*\**p* < 0.0001, one-way ANOVA. (**D-F**) mEPSCs recorded from control (*n* = 21) and KIF2C(WT) (*n* = 21) of cKO CA1 pyramidal neurons. n.s. *p* > 0.05, \**p* < 0.05, Student’s *t*-test; For the cumulative probability curves, \*\*\**p* < 0.001, Kolmogorov-Smirnov test. (**G**) Time course of percentage changes of fEPSCs amplitudes in cKO + control and cKO+KIF2C(WT) CA1 pyramidal neurons. cKO + control, *n* = 9 slices from 6 mice; cKO+KIF2C(WT), *n* = 8 slices from 6 mice. (**H**) Histogram shows fEPSC slope of cKO + control and cKO + KIF2C(WT) neurons (55-60 min). \*\**p* < 0.01. (**I-K**) mEPSCs recorded from cKO + control (*n* = 18) and cKO + KIF2C(G491A) (*n* = 15) CA1 pyramidal neurons. n.s. *p* > 0.05, Student’s *t*-test; For the cumulative probability curves, n.s. *p* > 0.05, \*\*\**p* < 0.001, Kolmogorov-Smirnov test. (**L**) Time course of percentage changes of fEPSCs amplitudes in cKO + control and cKO + KIF2C(G491A) CA1 pyramidal neurons. cKO, *n* = 7 slices from 6 mice; KIF2C(G491A), *n* = 8 slices from 6 mice. (**M**) Histogram shows fEPSC slope of cKO + control and cKO + KIF2C(G491A) neurons (55-60 min). n.s., *p* = 0.1791, Student’s *t*-test.

MTs remain dynamic in mature neurons and are capable of invading dendritic protrusions. MT invasion of dendritic protrusions depends on neuronal activity(*37*). To observe MT invasion, we used 50 ng/mL brain-derived neurotrophic factor (BDNF) to induce spine plasticity, and labeled newly polymerized MTs with an antibody against tyrosinated-α-tubulin. In the basal state, invading MTs were present more in the dendritic spines of cKO neurons (Figure 6C), suggesting increased MT invasion events in cKO synapses. After BDNF treatment, MT invasion robustly increased in WT neurons; however, no change was observed in cKO (Figure 6C). Based on our previous result showing KIF2C translocation from synapses to dendrites upon neuronal activity (Figure 2F–I), these results indicate that KIF2C is critical for activity-dependent microtubule invasion of spines. Synapses deficient in KIF2C have a moderate increase in basal microtubule invasion; however, they lose their ability to respond to neuronal activities due to the lack of further robust MT dynamics.

In mitotic cells, KIF2C binds to the end of MTs and couples ATP hydrolysis to initiate MT depolymerization(*38*). To further confirm whether the MT depolymerization property of KIF2C is critical for regulating synaptic transmission and plasticity, we constructed an MT depolymerization loss-of-function KIF2C mutant (KIF2C G491A). Its human homologous KIF2C G495A mutant has been previously reported to lose ATP hydrolyzing and hence MT depolymerization abilities (*20, 39*). The KIF2C(G491A) mutant was first tested in HEK 293T cells for its MT depolymerization function. Immunocytochemical analysis showed that KIF2C(G491A) transfected cells had intact cell morphology, while the KIF2C(WT) successfully depolymerized MTs and maintained cell rounding (Figure S3A). Furthermore, we expressed KIF2C(WT) or mutant in the hippocampal CA1 region of cKO mice by AAV-mediated gene delivery (Figure S3B, C) and examined synaptic transmission and plasticity using an electrophysiological approach. The results showed that re-expression of KIF2C(WT) significantly reduced the elevated amplitude of mEPSCs (Figure 6D–F) and restored LTP in cKO mice (Figure 6G, H). However, without MT depolymerization activity, the KIF2C (G491A) mutant failed to show significant differences in mEPSC properties (Figure 6I–K) or LTP induction in the cKO CA1 neurons (Figure 6L, M). These findings indicate that KIF2C regulates synaptic transmission and plasticity by mediating dynamic microtubule invasion of dendritic spines.

### KIF2C plays an important role in CA1-dependent cognitive behaviors

KIF2C deficiency leads to abnormalities in synaptic transmission and synaptic plasticity; we speculated whether KIF2C plays a role in higher-order brain function. Therefore, we examined KIF2C cKO mice for several behavioral tests. The KIF2C cKO mice showed normal motion and anxiety levels in the open field test and elevated zero maze compared to those of the WT mice (Figure S4A, B), and had normal grooming frequencies in the commonly used repetitive behavioral test (Figure S4C). The Morris water maze task also elicited similar learning curves and probe performance in both KIF2C cKO and WT mice (Figure S4D), suggesting normal spatial learning and memory ability in mice with KIF2C deficiency. However, when we subjected the mice to the Y-maze test, KIF2C cKO mice showed significant impairment in spontaneous alternation behavior compared to that of the WT mice (Figure 7A), indicating that KIF2C cKO mice were deficient in short working memory, which could be due to the special role of CA1 in temporal processing (*40-42*).

**Figure 7.**
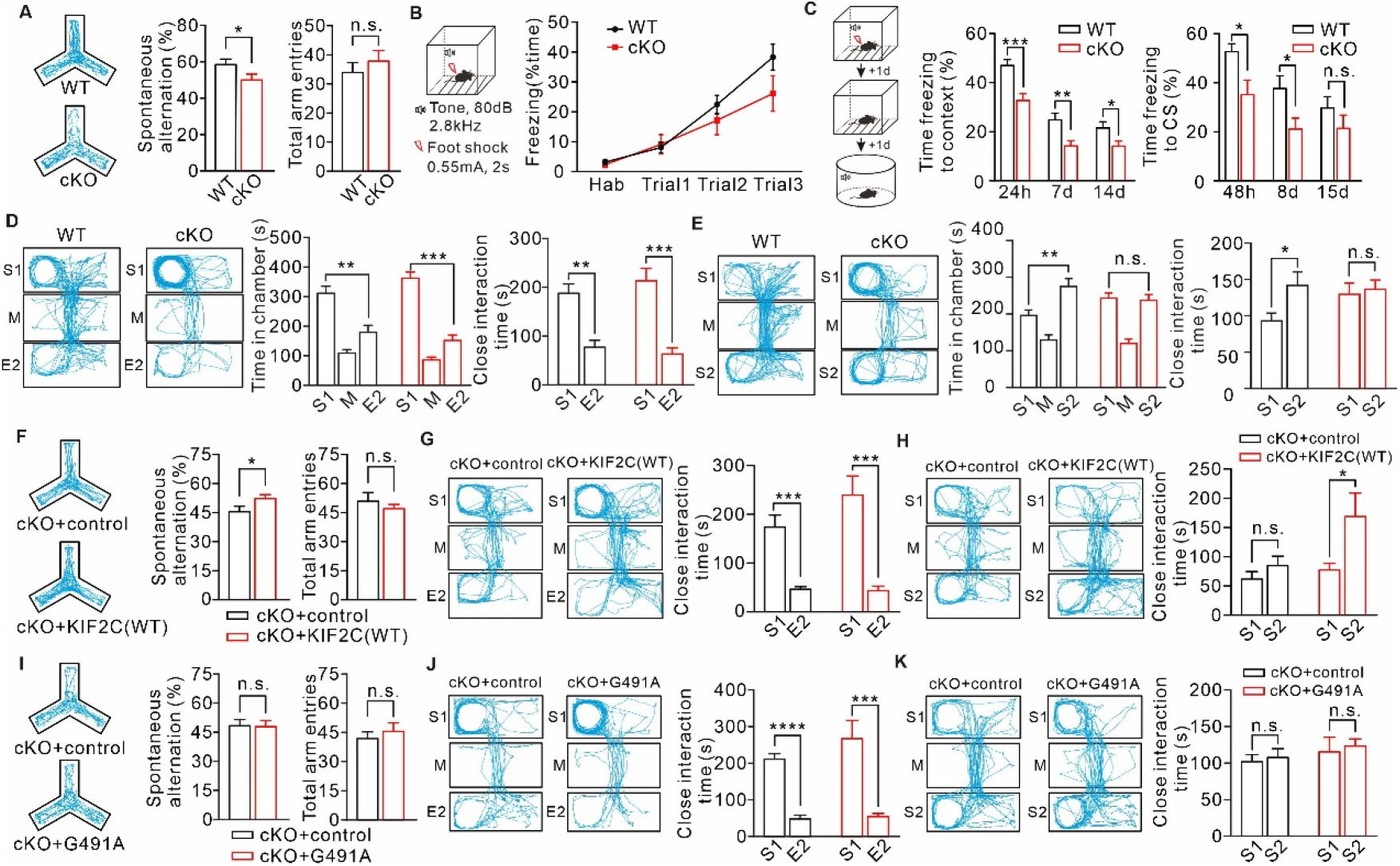
KIF2C cKO mice exhibit abnormal cognitive behaviors. (A) Y maze test from WT and cKO. *n* = 13 per genotype. n.s., *p* > 0.05, \**p* < 0.05, Student’s *t*-test. (**B-C**) Cued fear conditioning test. **B**, WT and cKO mice showed no significant freezing response to the tone CS across three CS-US pairings. **C**, Expression of conditioned freezing response to the background context or CS at indicated time. WT: *n* = 12; cKO: *n* = 10. Training, *p* > 0.05, two-way ANOVA. n.s., *p* > 0.05; \**p* < 0.05; \*\**p* < 0.01; \*\*\**p* < 0.001, Student’s *t*-test. (**D-E**) Three-chamber test. **D**, WT and cKO mice spent more time with stranger 1 in sociability test. \*\**p* < 0.01; \*\*\**p* < 0.001. **E**, cKO mice did not display a preference for stranger 2 in social novelty test. *n* =11 per genotype. n.s., *p* > 0.05, \*\**p* < 0.01, Student’s *t*-test. (**F**) cKO mice with KIF2C(WT) overexpression displayed a normal alternation in the Y-maze test. cKO+control, *n* = 9; cKO+KIF2C(WT), *n* = 7. n.s., *p* > 0.05, \**p* < 0.05, Student’s *t*-test. (**G-H**) KIF2C(WT) re-expression in CA1 can restore social novelty in cKO mice. cKO+control: *n* = 9; cKO+KIF2C(WT): *n* = 7. \**p* < 0.05, \*\*\**p* < 0.001. (**I**) cKO mice with KIF2C(G491A) overexpression displayed similar deficiency in the Y-maze test. n.s., *p* > 0.05, Student’s *t*-test. *n* = 13 per group. (**J-K**) cKO mice with KIF2C(G491A) overexpression displayed similar deficiency in social novelty test. *n* = 13 per group. n.s., *p* > 0.05, \*\*\**p* < 0.001, \*\*\*\**p* < 0.0001. Student’s *t*-test.

To investigate whether this learning deficit would translate to other cognitive disorders, we examined the mice for cued conditioned fear memory and social memory. The conditioned fear learning test showed that KIF2C cKO mice displayed a comparatively weaker freezing response to the context and tone conditioned stimulus (CS) during the acquisition stage (Figure 7B). We monitored the conditioned freezing response to the background context and CS until 14 or 15 days after acquisition, respectively, and found that cKO mice still showed less freezing compared to WT mice, particularly in context-dependent freezing (Figure 7C). These results suggest that the maintenance of fear memory in KIF2C cKO mice was impaired. In a prepulse inhibition test, WT and cKO mice showed the same extent of inhibition as the prepulse (Figure S4E), showing that the cKO mice had normal sensorimotor gating. In the three-chambered social test, both WT and KIF2C cKO mice exhibited no position preference in habituation stage (Figure S4F). KIF2C cKO mice exhibited a normal preference for the chamber in which a stranger mouse (S1) was present, compared with the empty chamber (Figure 7D). However, with the introduction of a second stranger (S2), KIF2C cKO mice exhibited no preference for the novel stranger (S2), compared with the familiar mouse (S1) (Figure 7E). These results showed that KIF2C cKO mice showed normal sociability but significantly reduced social novelty, indicating that KIF2C is essential for social memory maintenance.

Based on the behavioral tests, we found that KIF2C plays an important role in CA1-mediated cognitive and memory functions. The deficiencies observed in the Y-maze test, cued conditioned fear memory, and three-chambered social test of KIF2C cKO mice were highly likely due to the lack of LTP in the CA1 region (Figure 5G, H). As re-expression of KIF2C in the CA1 region restored LTP (Figure 6G, H), we speculated whether such CA1-specific rescue could also ameliorate the behavioral abnormalities in KIF2C cKO mice. In our pretests, over-expression of KIF2C in mice for a long time would lead to neuronal death (data not shown). Therefore, we re-expressed KIF2C in the cKO mice CA1 region using a doxycycline-induced viral expression system (Figure S4G) and examined the rescuing effects in the Y-maze test and three-chambered social test. The results showed that cKO mice with KIF2C(WT) overexpression displayed a normal alternation in the Y-maze test compared to cKO mice with control virus (Figure 7F). Furthermore, cKO mice with KIF2C(WT) overexpression also showed normal interaction with the stranger mouse in the social novelty test (Figure 7G, H). In addition, we tested whether the expression of the functional-null mutant KIF2C(G491A) in the CA1 region of cKO mice would have any rescue effect. The results showed that cKO mice injected with KIF2C(G491A) virus still had impaired working and social memory (Figure 7I–K). Therefore, these results suggest that KIF2C is critical for cognitive function, and this functional role is mediated by its MT depolymerization ability in synapses during LTP.

## Discussion

KIF2C is an ATP-dependent microtubule depolymerase that regulates cell division in many aspects of mitosis, such as spindle assembly, MT dynamics, correct kinetochore-MT attachment, and chromosome positioning and segregation(*38*). However, its function in the CNS remains unclear. This study elucidated the functions of KIF2C in regulating spine morphology, synaptic transmission, and plasticity in hippocampal neurons and its critical role in CA1-mediated cognitive behaviors. We demonstrated that the mechanism mediated by KIF2C is highly correlated with its known function in MT depolymerization and MT invasion of dendritic spines. This study expands the understanding of the function of KIF2C in non-mitotic cells and also unveils its critical role in the CNS.

The stabilization of dendritic spines is an important structural basis for synaptic plasticity. We found that both acute knockdown or genetic deficiency of KIF2C eliminated LTP except LTD expression. On a molecular basis, KIF2C in knockdown or cKO hippocampal CA1 neurons exhibited a higher AMPAR-mediated mEPSC amplitude and an increase in surface AMPARs expression, with a trend of increase in NMDARs. It is possible that increased expression of ionic glutamate receptors, particularly AMPARs, on synapses leads to LTP occlusion. It would be interesting to further explore the mechanism of KIF2C-involved receptor trafficking or synaptic expression. As important architectural elements in neurons, MTs are also responsible for intracellular transport. It has been reported that NMDA receptors can be transported by KIF17 (*43*) and AMPA receptors can be transported by KIF5 along MTs in dendrites (*44*). It is possible that KIF2C regulates membrane receptor trafficking by mediating MT dynamics through the regulation of MT stability. We observed microtubule dynamics using live cell imaging of KIF2C cKO hippocampal neurons. We observed that depletion of KIF2C induced faster, shorter-lived microtubule growth in dendrites. KIF2C deficiency also increased the frequency of MT invasion under basal conditions, which was probably due to the loss of MT polymerization suppression. Therefore, the increased AMPA receptor expression might be due to increased MT invasion, which brings AMPA receptors into synapses (Figure 8). Another hypothesis is that such changes in MT polymerization kinetics might affect the stability of AMPA receptor-containing vesicles on MT structures (Figure 8). It might cause the release of vesicles from MTs in the dendrite shaft, which may be distributed into spines by membrane infusion and later diffusion, similar to a previous report on the MT-dependent trafficking and synaptic entry of synaptotagmin-IV (*45*).

**Figure 8.**
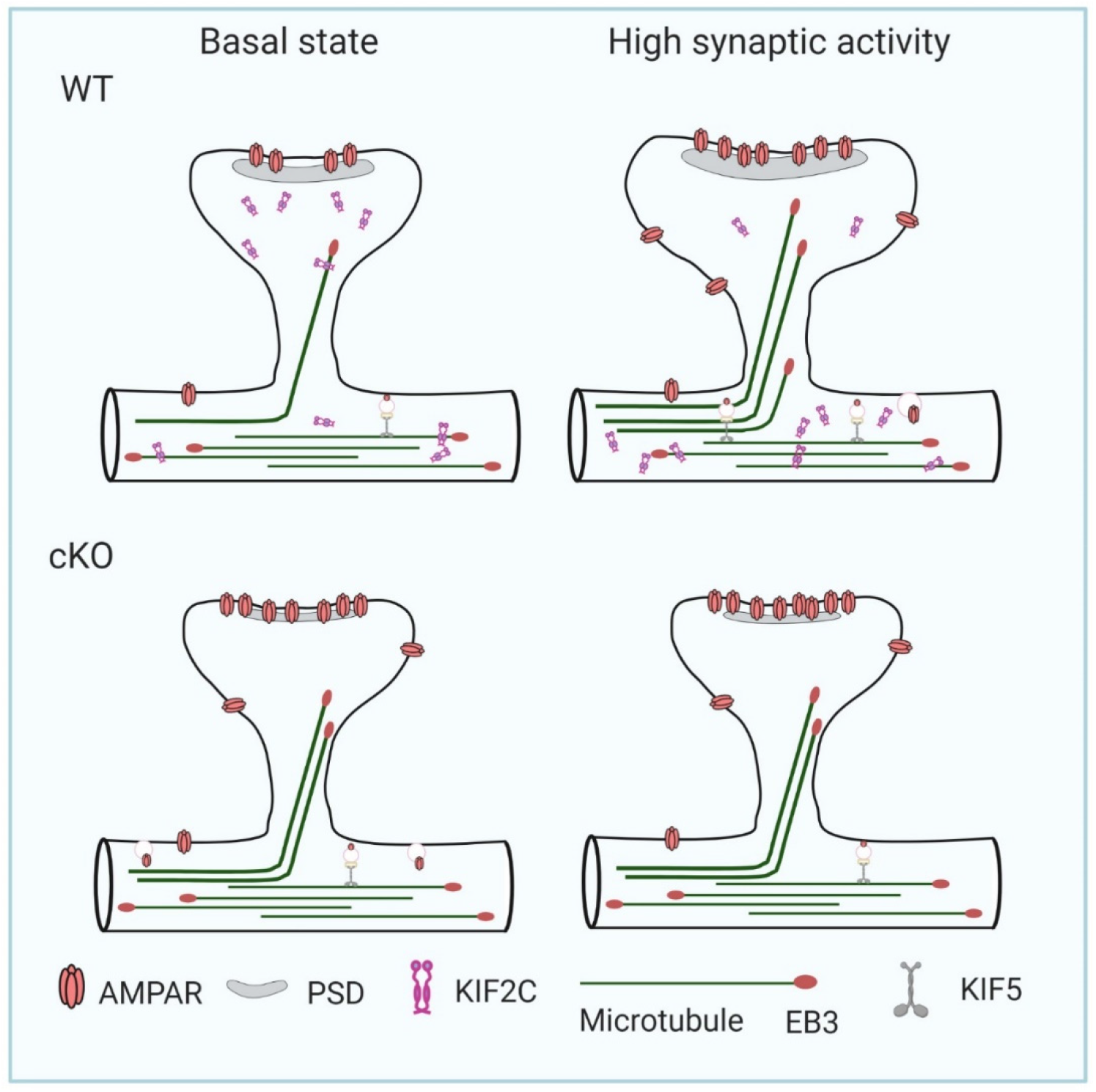
Model of KIF2C govern synaptic receptor expression by regulating dynamic MT synaptic invasion. KIF2C acts as a microtubule destabilizer in hippocampal neurons and highly expresses in the synapse. In basal state, KIF2C depolymerize MT to prevent excessive MT invasion into spine. During elevated synaptic activity, KIF2C translocate from synapse to dendritic shafts for more stable MT invasion into synapse in WT hippocampal neurons. In cKO neurons, loss of KIF2C leads to increased AMPA receptors membrane expression and the probability of MT invasion, impairs synaptic structure plasticity and destroy MTs activity-dependent invasion.

Activity-dependent MT invasion of dendritic spines becomes disabled after BDNF stimulation in cKO hippocampal neurons. Our findings reveal that during cLTP induction KIF2C was removed from the synapses (Figure 2F). Although the underlying mechanism is unclear, translocation of KIF2C from synapse to dendrite or depletion of synaptic KIF2C during synaptic activation is found to be required to ensure robust MT invasion into spines and where and when MT depolymerization events should be inhibited. Even though the genetic depletion of KIF2C increases the basal MT invasion into spines, it largely prevents further increase in MT invasion during LTP, resulting in synaptic plasticity failure and, finally, cognitive inflexibility (Figure 8). This hypothesis was further supported by our rescue experiments using the KIF2C (G491A) mutant, in which ATP hydrolyzing and MT depolymerization capabilities are abolished(*20*). Re-expression of KIF2C (WT), except for KIF2C (G491A) mutant, rescued the mEPSC hyper-amplitude in KIF2C cKO mice. KIF2C (G491A) also failed to restore the impaired LTP or cognitive function in KIF2C cKO mice during working memory or social memory tests as in the WT mice. Therefore, the MT depolymerization function of KIF2C plays a critical role in regulating synaptic plasticity and cognitive behaviors.

Interestingly, Oligomeric amyloid-β directly inhibits the MT-dependent ATPase activity of KIF2C *in vitro*, and thereby leads to the generation of defective mitotic structures which may contributes to the development of Alzheimer’s disease (*23*). Aβ stabilizes MT by reducing catastrophe-frequency (*46*), which is similar to KIF2C conditional knockout phenotype. Furthermore, KIF2C was recently discovered as a blood biomarker for suicidal ideation in psychiatric patients, such as patients with schizophrenia (*24*).Therefore, KIF2C could be a therapeutic target for Alzheimer’s disease or suicidality psychiatric disorders. Nevertheless, whether KIF2C plays other important role in higher-order brain functions remain to be explored.

## Materials and Methods

### Generation of KIF2C ^flox/flox^ mice and animal maintenance

KIF2C^flox/flox^ mouse was generated via CRISPR/Cas9 system in the Model Animal Research Center of Nanjing University (Nanjing, China). In brief, Cas9 mRNA, sgRNA and donor were co-injected into zygotes. sgRNA-5’ end: 5’-TCC ACT GTG GAA TGG TCT AA -3’; sgRNA-3’ end: 5’-CTG GTT GTG AGA CAC CTG TA-3’. Mouse *kif2c* gene is composed of 20 exons. The sgRNA directs Cas9 endonuclease cleavage in intron 1-2 and intron 8-9 and create a DBS (double-strand break). DBS breaks were repaired by homologous recombination, and resulted in LoxP sites inserted in intron 1-2 and intron 8-9 respectively. The mouse pups were genotyped by PCR and Southern blot followed by sequence analysis. Conditional knockout mice (KIF2C^flox/flox^; Nestin-Cre) were obtained by crossing KIF2C^flox/flox^ mice with Nestin-Cre mice (Wang et al., 2019). The resulting offspring were genotyped by PCR of genomic DNA. The primers information: KIF2C floxP fragment, F: 5’-GGT CCA GCT CTT TAC TGA TGT GTT C -3’; R: 5’-ACA AAG CAA GTC CAG GTC CAA G-3’; Nestin-Cre, F: 5’-TGC AAC GAG TGA TGA GGT TC-3’; R: 5’-GCT TGC ATG ATC TCC GGT AT-3’. Mice were kept under temperature-controlled conditions on a 12:12 h light/dark cycle with sufficient food and water *ad libitum*. Experiments were done in male littermates at the age of 1∼3 month. All animal experiments were performed in accordance with the ethical guidelines of the Zhejiang University Animal Experimentation Committee, and were in complete compliance with the National Institutes of Health Guide for the Care and Use of Laboratory Animals.

### Antibodies and reagents

Antibody to KIF2C (12139-1-AP, RRID: AB_2877829) was purchased from Proteintech (Rosemont, IL). The anti-PSD95 (ab2723, RRID: AB_303248), anti-MAP2 (ab5392, RRID: AB_2138153) and anti-EB3 (ab157217, RRID: AB_2890656) were purchased from Abcam (Cambridge, UK). The anti-α-tubulin, tyrosinated (MAB1864-I, RRID: AB_2890657), anti-synaptophysin (MAB368, RRID: AB_94947), anti-GAPDH (MAB374, RRID: AB_2107445), anti-GluA1(04-855, RRID: AB_1977216) and anti-GluA2 (MAB397, RRID: AB_2113875) were purchased from Millipore (Massachusetts, USA). The anti-GluN2A (4205, RRID: AB_2112295), anti-GluN2B (4207, RRID: AB_1264223), anti-GluN1 (5704, RRID: AB_1904067) were from Cell Signaling Technology (Danvers, MA). The anti-flotillin was from BD (610820, RRID: AB_398139). The anti-β-tubulin (sc-5274, RRID: AB_2288090) was from Santa Cruz (Dallas, TX). The anti-bassoon was from Enzo Life Sciences (ADI-VAM-PS003-F, RRID: AB_11181058). The phalloidin was from Yeasen (40734ES75, Shanghai, China). Horseradish peroxidase-conjugated secondary antibodies for immunoblotting were from Jackson and for immunostaining were from Invitrogen (Carlsbad, CA). Dulbecco’s modified Eagle’s medium (DMEM), Neurobasal medium, B27 were from Gibco. GlutMax, 4’,6-diamidino-2-phenylindole (DAPI), and Alexa Fluor-conjugated secondary antibodies were from Invitrogen (Carlsbad, CA). Nissl was from Beyotime (Shanghai, China). Protease inhibitor cocktail was from Merck Chemicals. Other chemicals were from Sigma (St. Louis, MO) unless stated otherwise.

### qRT-PCR

For single-cell analysis, the tip of a conventional patch-clamp pipette was placed tightly on the soma of a selected individual CA1 neuron. Gentle suction was applied to the pipette. After complete incorporation of the soma, the negative pressure was released and the pipette was quickly removed from the bath. The harvested contents were subjected to qRT-PCR using OneStep Kit (Qiagen, Hilden, Germany).

For tissue analysis, the hippocampus tissue RNA was extract by using RNeasy^®^ Mini Kit (Qiagen, Hilden, Germany). In brief, hippocampus tissue was homogenized in buffer RLT. The lysate was centrifuged at 12,000× g (4°C for 3 min). The supernatant was transferred and 1 volume of 70% ethanol was added. Then RNA was absorbed on RNeasy^®^ Mini spin column and purified by washing with buffer RW1 and RPE. The harvested contents were subjected to Reverse Transcription Kit (Vazyme). The qPCR reaction was performed on HiScript^®^ II 1st Strand cDNA Synthesis Kit (R212-02) (Vazyme, Nanjing, China) using ChamQ™ Universal SYBR^®^ qPCR Master Mix (Vazyme, Q711). Primer information: GAPDH: F 5’-AGG TCG GTG TGA ACG GAT TTG-3’; R 5’-TGT AGA CCA TGT AGT TGA GGT CA-3’. KIF2C: F 5’-TGC CGT TGT TGA TGG TCA GTG-3’; R 5’-GGA GAC ACT TGC TGG GAA CAG-3’. shKIF2C: F 5’-TGG ATC GAA GGA GGT ACC AC-3’; R 5’-CAC TGA CCA TCA ACA ACG GCA-3’. The following parameters were used: 95°C for 30s, followed by 39 cycles of 95°C for 10s, 60°C for 30s in accordance with the manufacturer’s protocol.

### Hippocampal neuron culture and transfection

Primary cultures of hippocampal neurons were prepared from embryonic day 18 mice. The isolated hippocampi were dissociated with 2.5% trypsin at 37°C for 20 min. The neurons were quantified and centrifuged at 1,800 rpm for 6 min and then plated on the glass coverslips pre-coated with poly-L-lysine (Sigma). The culture medium was Neurobasal (Gibco) supplemented with 2% B27 (Gibco), GlutMax (Invitrogen) and 1% Penicillin-Streptomycin in a 5% CO_2_ atmosphere at 37°C. Neurons were transfected at DIV 7-10 using a modified calcium phosphate protocol(*47*). More than 10 days after transfection, neurons were directly subjected to live imaging under a Nikon A1R confocal scanning microscope.

### DNA constructs

The shRNA targeting sequence is designed against position 820-838 of mouse KIF2C (Gene ID: 73804) open reading frame (5’-CCGGATGATCAAAGAATTT-3’), using Bgl II/SalI restriction enzymes. KIF2C(WT) uses the Nhe I/BamH I restriction enzymes. KIF2C point mutations were replaced a G to A at 491 position, using the Nhe I/BamH I restriction enzymes. Plasmid synthesis and virus construction were completed by the company (OBIO, Shanghai, China). KIF2C shRNA was sub-cloned into AAV backbone pAKD-CMV-bGlobin-mcherry-H1-shRNA. KIF2C (WT) was sub-cloned into AAV backbone pAAV-CMV-MCS-3FLAG-P2A-mNeonGreen-CW3SL and rTTA-TRE3G-mcherry (BrainVTA, Wuhan, China). KIF2C (G491A) was sub-cloned into AAV backbone pAAV-CMV-MCS-3FLAG-P2A-mNeonGreen-CW3SL (OBIO, Shanghai, China).

### Adeno-associated virus microinjection

For virus microinjection, WT or KIF2C cKO mice were randomly allocated to experimental groups and processed. A small craniotomy was performed after anesthetizing the mouse. The flow rate (40 nl/min) was controlled by a micro-injector (World Precision Instruments). Virus was microinjected in both left and right hippocampal CA1 region (from bregma, 3 weeks: ML: 1.45 mm; AP: 1.8 mm; DV: 1.5 mm; 2 months: ML: 1.65 mm; AP: 1.82 mm; DV: 1.62 mm). For electrophysiology, 4– 6-week-old mice were subjected for experiments at least 3 days after injection. For KIF2C overexpression mice subjected for behavioral tests, mice were injected and bred until 2–3-month-old, then took water containing (40 μg/ml) doxycycline for 3 days to induce KIF2C expression before tests.

### Glycine-induced LTP

Hippocampal neurons were infected by shRNA virus at DIV 7-10, and subjected for chemical LTP induction at DIV 16-18. Briefly, neurons were firstly incubated in extracellular solution (ECS; in mM: 140 NaCl, 2 CaCl_2_, 5 KCl, 5 HEPES, 20 glucose) supplemented with 500 nM TTX, 1 μM strychnine, 20 μM bicuculline (Buffer A; pH 7.4) at 37°C for 5 min, and treated with 200 μM glycine in Buffer A for 10 min. Then neurons were incubated in Buffer A for another 5-20 min. The cells were then harvested for experiments(*32, 48, 49*).

### Western blots

After determining protein concentration with BCA protein assay (Bio-Rad), equal quantities of proteins were loaded and fractionated on SDS-PAGE and transferred to NT nitrocellulose membrane (Pall Corporation), immunoblotted with antibodies, and visualized by enhanced chemiluminescence (Pierce Biotechnology). Primary antibody dilutions used were GluA1 (1:1,000), GluA2 (1:2,000), GluN1 (1:2,000), GluN2A (1:2,000), GluN2B (1:2,000), flotillin (1:1,000), KIF2C (1:500), EB3 (1:10,000), PSD95 (1:1,000), β-tubulin (1:2,000), GAPDH (1:10,000), and secondary antibodies (1:10,000). Film signals were digitally scanned and quantitated using Image J software.

### Immunocytochemistry

DIV 18-20 hippocampal neurons were fixed in 4% paraformaldehyde (PFA) containing 4% sucrose for 20 min, and then treated with 1% Triton X-100 in PBS for 10 min at room temperature (RT). After washing with 1x PBS, neurons were transferred into blocking solution (10% normal donkey serum, NDS in PBS) for 1 h at RT. Neurons were then incubated with primary antibodies at 4°C overnight and incubated with secondary antibodies for 3 h at RT. The samples were subjected to confocal laser scanning microscope Nikon A1R. Primary antibody dilutions used for immunocytochemistry were PSD95 (1:1,000), KIF2C (1:500), bassoon (1:1,000), tyrosinated α-tubulin (1:500), and MAP2 (1:10,000), and secondary antibodies (1:1,000). All antibodies were diluted in 1x PBS containing 3% NDS. Measures of spine head width and spine density in cultured hippocampal neurons were performed using Imaris software and colocalization analysis was performed by Metamorph software.

### Immunohistochemistry

Mice were anesthetized and transcranially perfused with 4% PFA. Coronal slices with 30 μm thickness were prepared and placed in blocking solution for 1 h at RT. After washing with 1x PBS, slices were incubated with primary antibodies overnight at 4°C and incubated with secondary antibodies for 3 h at RT. Images were acquired using an automated fluorescence microscope Olympus BX53.

### Nissl staining

Nissl staining was performed using Nissl staining Kit (Beyotime, C0117). Coronal slices (30 μM thickness) were immersed in Nissl staining solution for 10 min at 37°C, rinsed with ddH_2_O, dehydrated in ethanol, and cleared in xylene. Images of cortex and hippocampus were captured using an Olympus VS120 microscope. Hippocampus CA1 and CA3 thickness was calculated by Image J software.

### Golgi staining

Golgi-Cox staining of hippocampus CA1 tissue was performed using the FD Rapid GolgiStain™ Kit (FD NeuroTechnologies, PK401) following the manufacturer’s protocol. In brief, mice were deeply anesthetized with intraperitoneal injection of 4% chloral hydrate (10 μl/g), the brain was rapidly removed and rinsed briefly in ddH_2_O. The tissues were immersed in impregnation solution made by mixing equal volumes of solution A and B, and then stored at RT for 2 weeks in the dark. Tissues were transferred to solution C and store at 4°C in dark for 7 days. Coronal slices (150 µm thickness) were prepared using a vibrating tissue slicer (Leica VT1000S) in solution C. Slices were mounted on gelatin-coated slides (FD NeuroTechnologies, PO101) with solution C and dry naturally at RT. Slices were rinsed in ddH_2_O twice and placed in a mixture consisting of 1 part solution D, 1 part solution E and 2 part ddH_2_O for 10 min. Slices were rinsed in ddH_2_O twice, dehydrated in 50%, 75%, 95% and 100% ethanol for 4 min, cleared with xylene for 3 times, sealed with permount (Sigma, 06522). Images of hippocampus CA1 pyramidal neurons were captured by an Olympus BX61 microscope with 100x objective lens. The sholl analysis, spine density and the morphological classification of the spines were calculated by Image J software(*50*). The morphological classification of spines as previously described(*34, 51, 52*): mushroom, big protrusions with a well-defined neck and a very voluminous head; stubby, short protrusions without a distinguishable neck and head; long-thin, protrusions with a long neck and a clearly visible small bulbous head; filopodia, thin, hair-like protrusions without bulbous head.

### Preparation of subcellular fractionation

Hippocampal tissues were homogenized in buffer C (320 mM sucrose, 20 mM Tris-HCl [pH=8.0], 2 mM EDTA and 200 μg/ml PMSF) supplemented with protease inhibitors (Merck Chemicals) and centrifuged at 800x g for 10 min at 4°C to result in P1 and supernatant S1. The supernatant S1 was centrifugated at 12,000x g for 15 min at 4°C to yield P2 and S2. Then P2 fraction was resuspended in buffer D (20 mM Tris-HCl [pH=8.0], 1 mM EDTA, 100 mM NaCl, and 1.3% Triton X-100) and centrifuged at 100,000x g for 1 h to produce a supernatant S3 and a pellet P3. The pellet represented PSD fraction, and the protein concentration was measured using BCA protein assay kit (Thermo, 23225).

### Surface biotinylation assay

Acute mouse coronal slices containing hippocampus region (250 μm) were prepared from anesthetic mice using a vibrating tissue slicer (Leica VT1000S) in ice-cold artificial cerebrospinal fluid (ACSF) containing: 125 mM NaCl, 2.5 mM KCl, 1.25 mM NaH_2_PO_4_, 1 mM MgCl_2_, 2 mM CaCl_2_, 26 mM NaHCO_3_ and 25 mM D-glucose, bubbled with 95% O_2_ / 5% CO_2_. After recovery for 30 min at 37°C, slices were incubated with 0.5 mg/ml EZ-link-sulfo-NHS-SS-Biotin (Thermo, 21331) in ACSF for 30 min. The biotin solution was then removed, and the remaining unreacted biotin was quenched by the addition of 50 mM Tris solution (pH=7.4) for 2 min. The slices were then washed with ice-cold PBS and lysed with RIPA buffer supplemented with protease inhibitors. Biotinylated proteins were separated from other proteins using streptavidin agarose beads (Thermo, 20349) overnight at 4°C. The agarose beads were then washed three times with RIPA buffer and 500× g, 2 min centrifuge. The biotinylated proteins were extracted with SDS-PAGE loading buffer and boiled for 3 min before western blot.

### Transmission Electron Microscopy (TEM)

Mice (P30) were transcranial perfused with saline and ice-cold fixative (4% PFA containing 0.1% glutaraldehyde), brains were removed and stored at 4°C for 2.5 h in fixative. Coronal slices containing hippocampus region (250 μm) were cut on a vibratome and kept in ice-cold fixative solution. Small blocks of tissue (1 mm^2^) from the CA1 region were excised under a microscope. The CA1 tissues were then rinsed for 6 times with 0.1 M PB and post-fixed in 1% OsO_4_ for 30 min, rinsed for 3 times with ddH_2_O and stained with 2% uranyl acetate for 30 min at room temperature. After dehydrating through a graded series of ethanol (50%, 70%, 90% and 100%), the samples were embedded in an epoxy resin. Ultrathin sections (90 nm) were cut using an ultra-microtome (Leica), stained with lead citrate solution, and mounted on grids. EM images were captured at 11,000× and 68,000× magnification using a Tecnai transmission electron microscope (FEI, Hillsboro, OR). CA1 synapses were identified by the presence of a presynaptic bouton with at least two synaptic vesicles within a 50 nm distance from the cellular membrane facing the spine, a visible synaptic cleft, and PSD(*53*). Image J software was used to analyze the PSD length and PSD thickness.

### Electrophysiology

We used 1% barbital sodium to anesthetize mice and removed the brain of mice rapidly and place it in ice-cold, high-sucrose cutting solution containing (in mM): 194 Sucrose, 30 NaCl, 4.5 KCl, 0.2 CaCl_2_, 2 MgCl_2_, 1.2 NaH_2_PO_4_, 26 NaHCO_3_, 10 Glucose. We cut slices coronally in the high-sucrose cutting solution with vibratome, and transferred the slices to an incubation chamber with artificial cerebrospinal fluid (ACSF) containing (in mM): 119 NaCl, 2.5 KCl, 2.5 CaCl_2_, 1.3 MgSO_4_, 1.0 NaH_2_PO_4_, 26.2 NaHCO_3_, 11 Glucose. Slices were incubated at 34 °C for 30 min and then transferred to 27°C for at least 1 hour before recording. During recording, the slices were placed in a recording chamber constantly perfused with warmed ACSF(32°C)and gassed continuously with 95% O_2_ and 5% CO_2_. Pipettes for Whole-cell recordings (3-5 MΩ) were filled with a solution containing (in mM): 115 CsMeSO_4_, 20 CsCl, 10 HEPES, 2.5 MgCl_2_, 4 Na-ATP, 0.4 Na_2_-GTP, 10 Na_2_phosphocreatine, 0.6 EGTA, 5 QX-314 (pH 7.2-7.4, osmolarity 290–310). Data were collected with a MultiClamp 700B amplifier and analyzed by pClamp10 software (Molecular Devices, Sunnyvale, USA). The initial access resistance was <25 MΩ and was monitored throughout each experiment. Data were discarded if the access resistance changed 20% during whole recording. Data were filtered at 2 kHz and digitized at 10 kHz.

For field excitatory postsynaptic potential recording, DG zone was cut down before recording to avoid the generation of epileptic wave. A concentric bipolar stimulating electrode was placed in the stratum radiatum to evoke EPSP and the recording electrode filled with ACSF was placed at 200-400 µm away from the stimulus electrode at the same region. Increasing stimulus intensity step by step to find the maximal response and then the baseline response was adjusted at 1/3 to 1/2 of the maximal response. After the steady baseline response was recorded for 30 min, a strain of 100 Hz-100 pulses high frequency stimulus was delivered to induce long-term potential (LTP) and a train of 2 Hz-900 pulses low frequency stimulus was delivered to induce long-term depression (LTD). The induced post response was recorded for 1 hour. The slope within 2 ms at initial EPSP was calculated as the value for field EPSP. The magnitude of LTP and LTD was calculated based on the EPSP values 50-60 min after the end of the induction protocol.

For whole-cell LTP recording, 100 µmol/L picrotoxin was added in recycling ACSF to block GABAR-mediated current. The stimulus electrode was placed in the stratum radiatum to evoke EPSC at 0.1 HZ and AMPAR-mediated current was recorded at -70 mV. After the steady baseline response was recorded for 5 min, the cell was held at 0 mV and received 2 strains of 100 Hz-100 pulses high frequency stimulus separated by 20 s. And then the cell was held at -70 mV to record for 40 min. Here the HFS must be given within 10 min of achieving the whole-cell recording to avoid “wash-out” of LTP. The magnitude of LTP was calculated based on the EPSC values 35-40 min after the end of the induction protocol.

For NMDAR-mediated current and AMPAR-mediated current recording, 100 µmol/L picrotoxin was added in recycling ACSF to block GABAR-mediated current. AMPAR-mediated current was recorded at -70 mV. A mixed current for AMPAR-mediated current and NMDAR-mediated current was recorded at +40 mV. The current at 50 ms after stimulus was counted for NMDAR-mediated current. The AMPAR/NMDAR ratio was calculated as the peak of the averaged AMPAR-mediated EPSC (30-40 consecutive events) at -70 mV divided by the averaged NMDAR-mediated EPSC (30-40 consecutive events).

For mini excitatory synaptic transmission recording, 100 µmol/L picrotoxin and 1 µmol/L TTX was added into recycling ACSF to block GABA currents and sodium currents respectively. Recording was conducted at -70 mV. Mini analysis software was used to count the mini EPSCs.

Paired-pulse facilitation (PPF) was examined by paired stimulations at different intervals.

### Behavioral experiments

All behavioral experiments were performed with the age of 2-to 3-month-old male littermates. All animals were given at least 1 hours of habituation time before testing.

#### Open field test

The open field test was used to evaluate the autonomous behavior, exploratory behavior and tension of animals in a new environment. Through dynamic monitoring of exploratory behavior and spontaneous activity, it is often used to test basic exercise volume and anxiety level. The mice were put in an area of 40 cm× 40 cm× 40 cm box and exploring freely for 30 min using a video tracking system. Data were analyzed using ANY-MAZE software.

#### Y maze

The Y-maze was used to evaluate discriminative learning, short-term working memory and reference memory test. It is based on the innate curiosity of the rodents to explore an arm that has not been previously visited. The Y-shaped maze is composed of three equal-length arms (40 cm×9 cm×16 cm). The angle between each two arms is 120 degrees. The inner arms and the bottom of the maze are painted dark grey. The mice were placed in the center of the maze and allowed to freely explore the three arms for 8 min. Take the animals’ four feet into the arm as the standard. One alternation would be recorded if all arms were entered successively at a time. The numbers of arm entries were recorded to evaluate the locomotion of animals in the Y-maze. Spontaneous alternation was calculated as total alternations/the number of maximum alternations (total arm entries-2).

#### Elevated zero maze

Elevated zero maze was used to investigate the animal’s anxiety state which was based on the animal’s exploring characteristics of the new environment and the fear of hanging open arms to form conflicting behaviors. The annular platform (60 cm in diameter, 5 cm track width) was evaluated 50 cm above the floor and divided equally into four quadrants. Two opposite quadrants were “open” and others were surrounded by 16 cm high blue, opaque walls. The mice were randomly placed in closed quadrants and allowed to freely to explore for a period of 5 min using an overhead video tracking system. Dependent measures were: time in open quadrants, open entrances, number of head dips, number of stretching and number of four paws in open section.

#### Morris water maze

Morris water maze is the most classic behavioral experiment that evaluates animal learning, memory, cognition, and space exploration ability. The water maze consists of a circular pool (radius 120 cm) filed with water containing nontoxic titanium pigment to make the platform invisible. Mice were trained to find the hidden platform (5 cm) and placed gently in different quadrants each time. The maximum searching period allotted was 60s. If a mouse did not find the platform, it will be placed on the platform by the experimenter and remained for 10s. Training was performed for five days and tested on the sixth day. For the training sessions, the time taken to find the platform, velocity and distance traveled were recorded. For the test session, the platform was removed. The time spent in the four quadrants of the pool were recorded.

#### Self-grooming

The mice were placed individually in a clean cage with fresh bedding. After 10 min habituation, mice were monitored with a camera and recorded for 10 min to score the spontaneous grooming behavior. Time of grooming was recorded.

#### Three-chambered social test

Three-chamber social interaction test was carried out as described previously(*54*). Briefly, two empty steel cylinders were respectively placed in the left and right rectangular. The test mice were individually placed in the center chamber for a 10 min habituation. Then the mice encountered a stranger C57BL /6Nmouse (S1) in the cage for 10 min. The social novelty test was then arranged and another stranger C57BL/6N (S2) was placed in the cage and let the test mouse explored freely for 10 min. The close interaction time of the test mouse spend in each chamber was recorded.

#### Cued fear conditioning

Mice from each genotype were placed to the conditioning chamber for a 160s habituation and then received 3 CS-US trails inter-spaced by 40s inter-trail intervals. They were given 60s stimulus-free interval before returned to their home cage. The conditional stimulus (CS) comprised a 20s discrete tone (80 dB, 2.8 kHz), which was followed by an unconditional stimulus at the last 2s (US, 0.55 mA foot shock). To examine the condition response to the background contextual cues, the mice were placed back in the conditioning chamber for 6 min in the absence of the CS and US stimuli after training 24h, 7d and 14d. To examine the condition response to the CS, the mice were placed in a distinct and modified context and given a 6 min CS tone after training 48h, 8d and 15d. For all procedures, the animal freezing behaviors were monitored using a manufacturer’s software.

#### Prepulse inhibition (PPI) test

PPI was performed as described previously(*55*). Briefly, in PPI test, prepulses (PP) were set at 70,74,78 and 82 dB, and pulse (P) was set at 120 dB. Percentage PPI was calculated as [(P amplitude – PP amplitude)/ P amplitude] x 100%.

### Statistics

Data were analyzed using GraphPad Prism 8.0 (GraphPad Software, San Diego, CA), Excel 2017 (Microsoft, Seattle, WA), and sigmaplot 12.0 (Systat Software, Erkrath, Germany). Data analysts were blind to experimental conditions until the data were integrated. Standard deviations for controls were calculated from the average of all control data. Statistical difference was determined using two-sided unpaired Student’s *t* test or One-way ANOVA. The accepted level of significance was *p* < 0.05. *n* represents the number of preparations, experimental repeats or cells as indicated in figure legends. Data are presented as mean ± SEM.

## Acknowledgments

We are grateful to research assistant Shuang-shuang Liu from the Core Facilities of Zhejiang University School of Medicine, Drs. San-hua Fang and research assistant Daohui Zhang from the Core Facilities of Zhejiang University Institute of Neuroscience and research assistant Bei-bei Wang from the center of Cryo-Electron Microscopy of Zhejiang University.

## Author contributions

R.Z. and J.X. conceived and designed the study. R.Z. and Y.-L.D. performed experiments. R.Z., Y.-L.D., X.-T.W., T.-L.L., Z.Z., N.W., X.-M.L., Y.S., J.-H.L., J.X., Z.W. and J.X. analyzed and discussed the data. R.Z. and J.X. wrote the initial manuscript, Y.S., J.-H.L., J.X., Z.W., L.S. and J.X. revised the manuscript. All authors commented on the manuscript.

## Figures

**Supplementary Information Figure S1:**
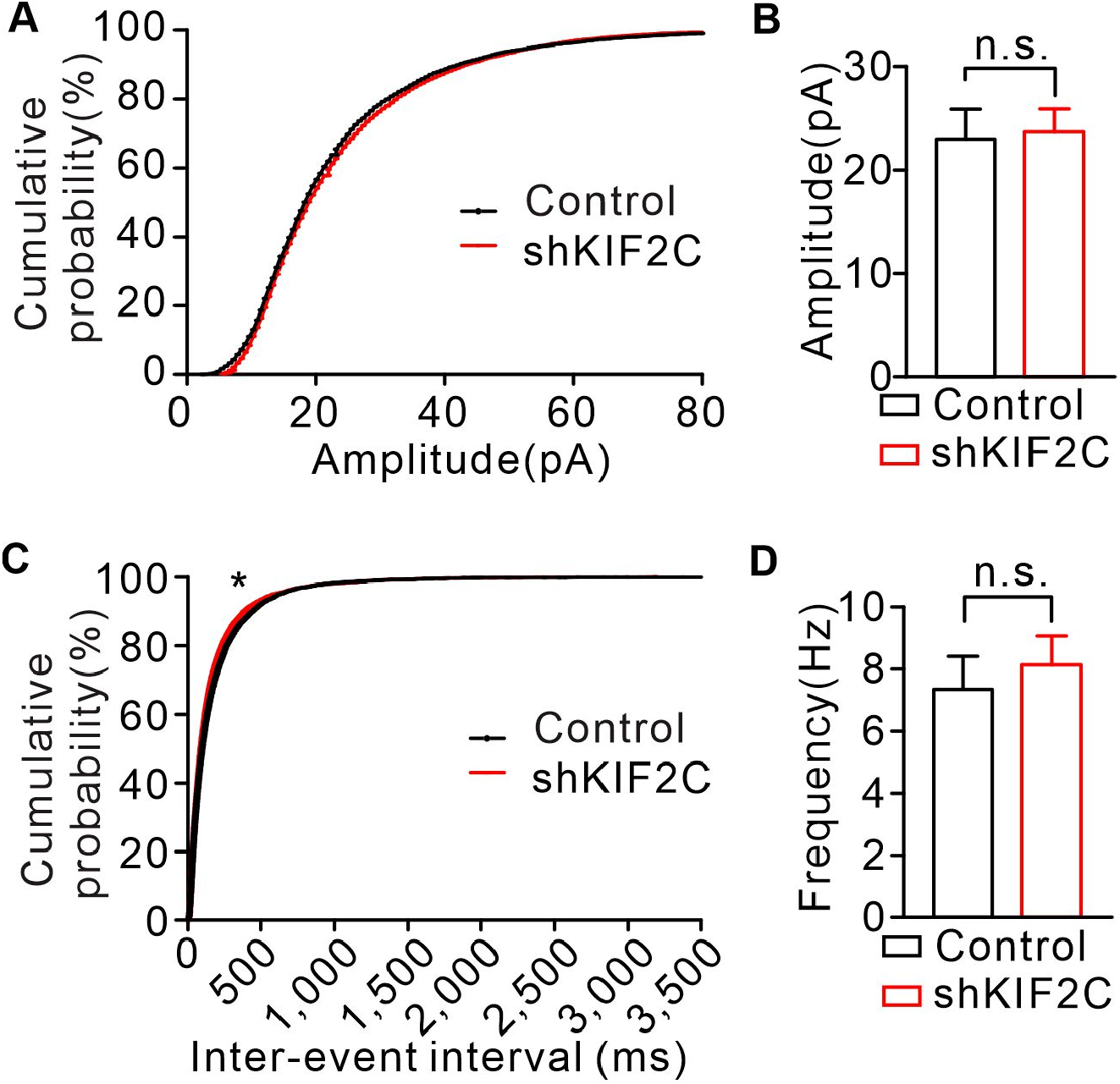
miniature excitatory synaptic currents (mEPSCs) recordings of cultured WT and KIF2C knockdown (shKIF2C) neurons. (A) mEPSCs recorded from WT (*n* = 14) and shKIF2C (*n* = 16) of cultured hippocampus neuron at DIV16-18. Averages of amplitude were 23.0 ± 2.76 pA (control) and 23.67 ± 2.45 pA (shKIF2C), *p* =0.7123, Student’s *t*-test; *p* > 0.05, Kolmogorov-Smirnov test. (**B**) Averages of frequency were 7.34 ± 1.02 Hz (control) and 8.13 ± 1.03 Hz (shKIF2C). *p* = 0.5828 Student’s *t*-test. \**p* < 0.05, Kolmogorov-Smirnov test.

**Supplementary Information Figure S2:**
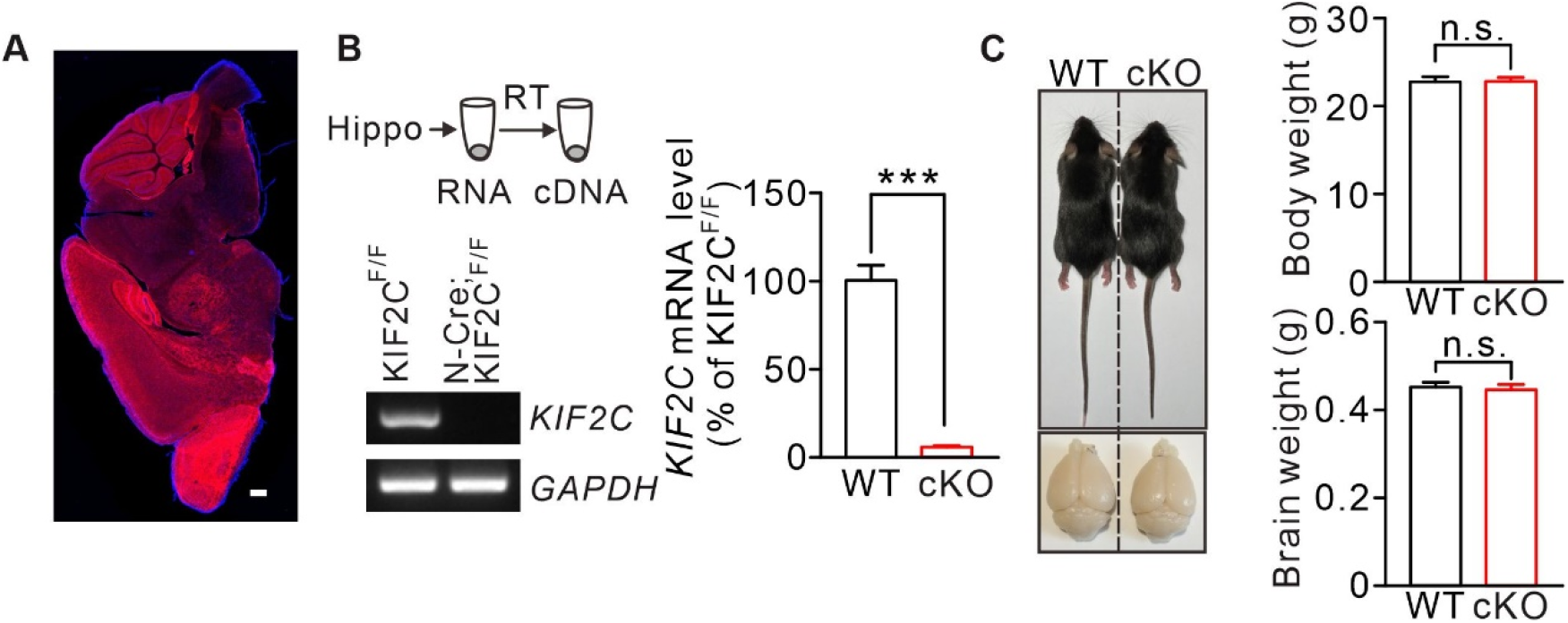
Nestin-Cre;KIF2C^flox/flox^ mice. (A) Nestin-Cre;KIF2C^flox/flox^ mouse was crossed with Ai9 reporter mouse lines and the expression of Cre-recombinase was characterized by observing tdTomato reporter in Nestin-Cre;Ai9 mice (1-month). We found that tdTomato fluorescence was present in the cortex, hippocampus, cerebellum and many other brain regions. Scale bar: 500 μm. (B) Electrophoresis of *KIF2C* (387 bp), and *GAPDH* (220 bp) amplicons from individual WT (*n* = 10) and cKO (*n* = 10) hippocampus tissues. Right histograms show percentage changes of *KIF2C* mRNA levels (WT: 100.0 ± 8.8%; cKO: 5.7 ± 0.9%, \*\*\**p* < 0.001, Student’s *t*-test). (**C**) cKO mice (2-month) displayed normal body weight and brain size. Body weight: WT, 22.77 ± 0.6 g; cKO, 22.8 ± 0.5 g, n = 10 per genotype. n.s., *p* > 0.05, Student’s *t*-test. Brain weight: WT, 0.45 ± 0.01 g; cKO, 0.44 ± 0.01 g, *n* = 8 per genotype. n.s., *p* > 0.05, Student’s *t*-test.

**Supplementary Information Figure S3:**
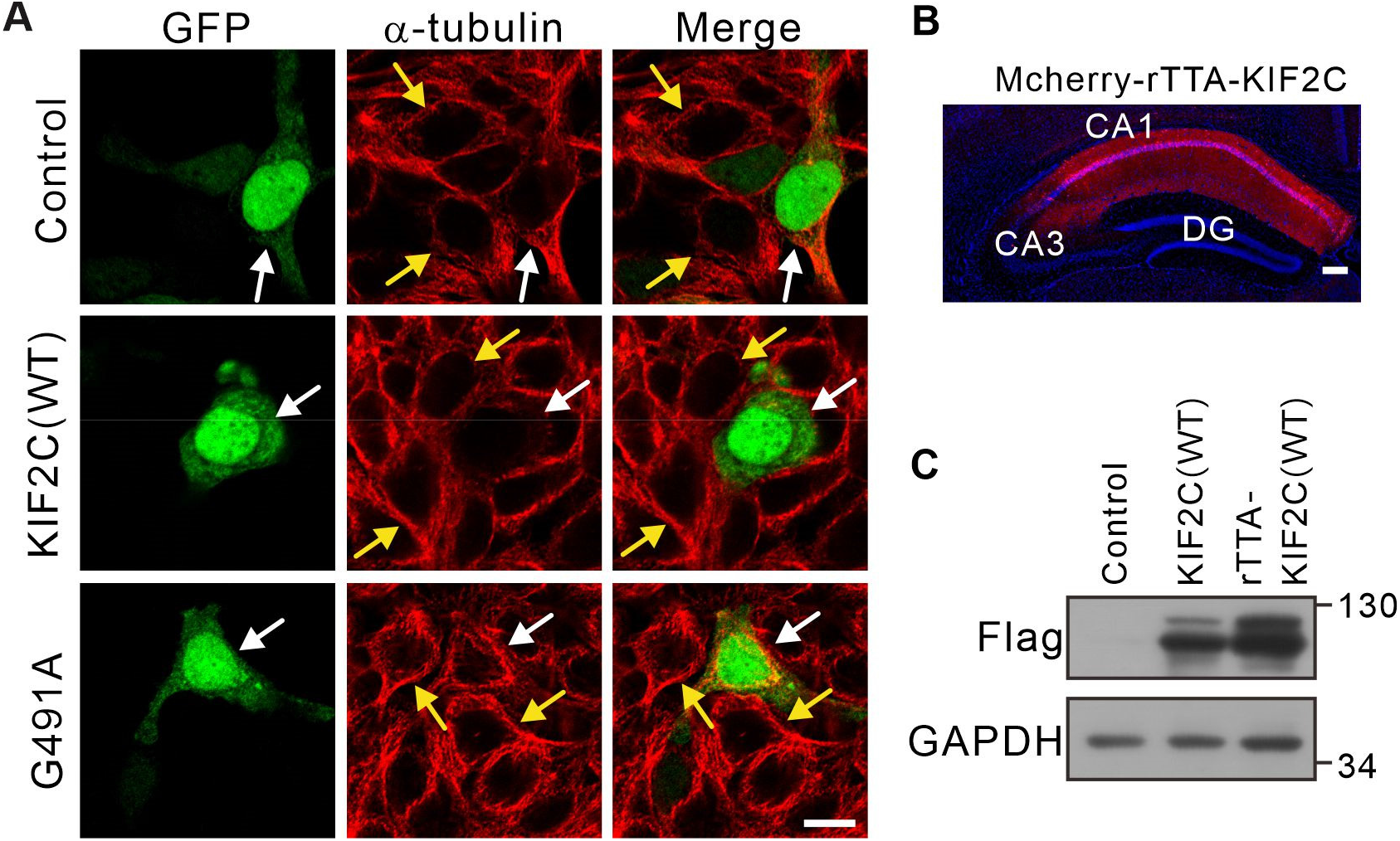
KIF2C(WT) and KIF2C(G491A) expression. (A) GFP vector, KIF2C(WT) and KIF2C(G491A) mutant were transfected into HEK 293T cells and α-tubulin was stained. White arrow points to transfected cells, yellow arrows point to intact cells. Scale bar: 10 μm. (**B**) Coronal section of the CA1 region image after mcherry-rAAV-KIF2C virus injection. (**C**) Expression of Flag-KIF2C (WT) in CA1 region total protein, GAPDH was the internal control.

**Supplementary Information Figure S4:**
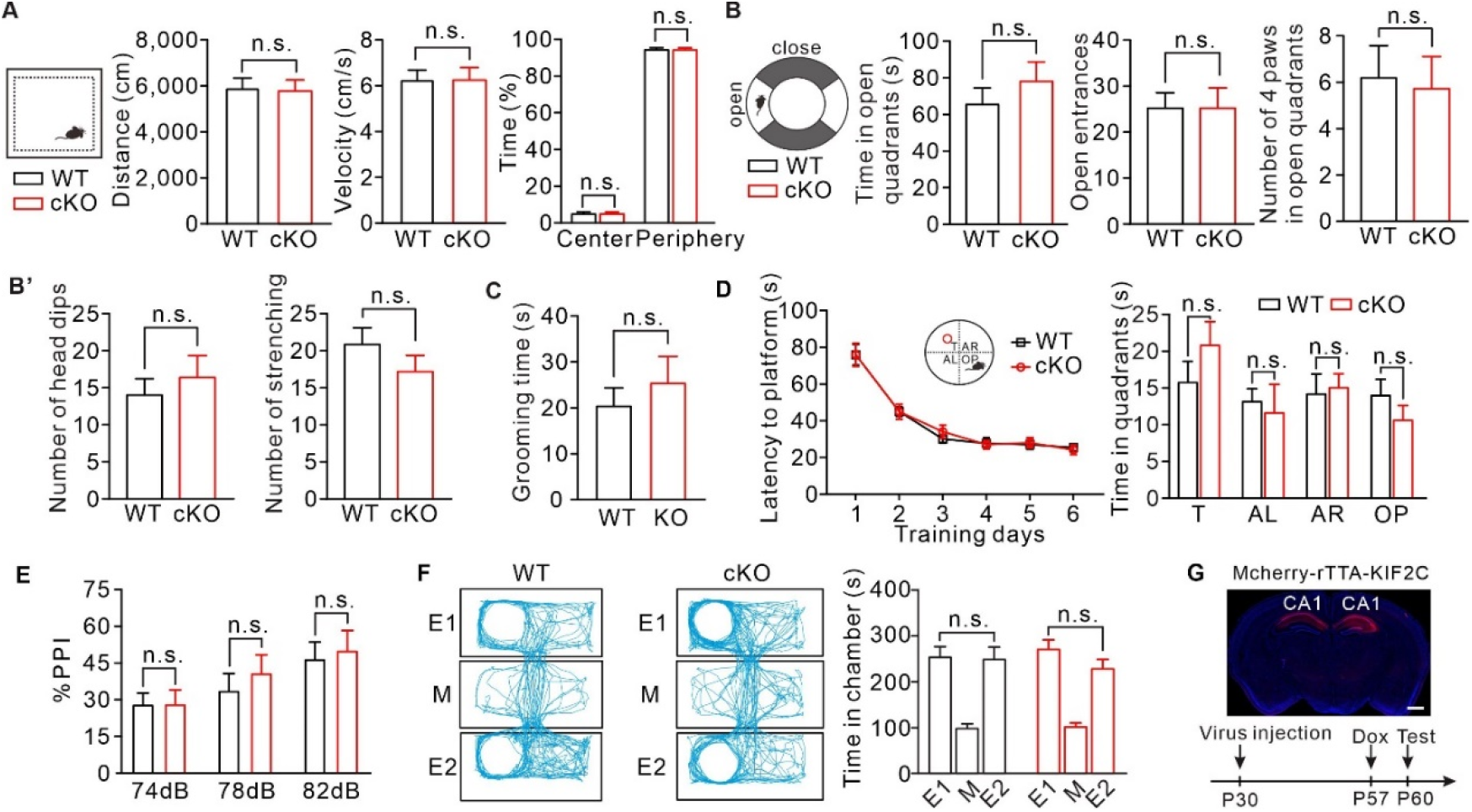
WT and KIF2C cKO mice behavioral tests. (**A**) Open field test from WT and cKO mice. Quantification of velocity, distance of activities and exploration time. *n* = 13 per genotype. (**B**) Elevated-zero maze test. *n* = 11 per genotype. (**C**) Self-grooming test. WT: *n* = 15; cKO: *n* = 18. (**D**) WT and cKO mice displayed similar latency to platform over training days in Morris water maze test. T, target; AL, adjacent left; AR, adjacent right; OP, opposite. n.s., *p* > 0.05; two-way ANOVA. (**E**) Prepulse inhibition (PPI) test. Pulse intensity was set to 120 dB. WT: *n* = 10; cKO: *n* = 9. Data are represented as means ± SEMs. n.s., *p* >0.05, Student’s *t*-test. (**F**) Both WT and KIF2C cKO mice show no position preference in habituation stage. n.s., *p* > 0.05. (**G**) Time line of virus injection, doxycycline (Dox) administration and behavior test. Coronal image of rAAV-TRE3G-mcherry-KIF2C re-expression in CA1. Scale bar: 800 μm.

